# A novel antiviral lncRNA EDAL shields a T309 *O*-GlcNAcylation site to promote EZH2 degradation

**DOI:** 10.1101/824813

**Authors:** Baokun Sui, Dong Chen, Wei Liu, Qiong Wu, Bin Tian, Jing Hou, Yingying Li, Shiyong Liu, Juan Xie, Hao Jiang, Zhaochen Luo, Lei Lv, Fei Huang, Ruiming Li, Min Cui, Ming Zhou, Huanchun Chen, Zhen F. Fu, Yi Zhang, Ling Zhao

## Abstract

The central nervous system (CNS) is vulnerable for viral infection, yet few host factors in the CNS are known to defend invasion by neurotropic viruses. We report here that multiple neurotropic viruses, including rabies virus (RABV), vesicular stomatitis virus (VSV), Semliki Forest virus (SFV) and herpes simplex virus 1 (HSV-1), elicit the neuronal expression of a host-encoded lncRNA EDAL. EDAL inhibits the replication of these neurotropic viruses in neuronal cells and RABV infection in mouse brains. EDAL binds to the conserved histone methyltransferase enhancer of zest homolog 2 (EZH2) and specifically causes EZH2 degradation via lysosomes, reducing the cellular H3K27me3 level. The antiviral function of EDAL resides in a 56-nt antiviral substructure through which its 18-nt helix-loop intimately contacts multiple EZH2 sites surrounding T309, a known *O*-GlcNAcylation site. EDAL positively regulate the transcription of *Pcp4l1* encoding a 10 kDa peptide, which inhibits the replication of mutiple neurotropic viruses. Our findings proposed a model in which a neuronal lncRNA can exert an effective antiviral function via blocking a specific *O*-GlcNAcylation that determines EZH2 lysosomal degradation.

## INTRODUCTION

Among infectious diseases of the central nervous system (CNS), those caused by viral pathogens—known as neurotropic viruses—are far more common than bacteria, fungi, and protozoans (2Nd & Mcgavern, 2015, Ludlow, Kortekaas et al., 2016). Neurotropic viruses arrive to the CNS through multiple routes and propagate within various cell types including astrocytes, microglia and neurons, depending on the entering routes and virus types (Manglani & McGavern, 2018). Infection of some neurotropic viruses can cause meningitis or encephalitis and result in severe neurologic dysfunction, such as VSV, SFV, HSV-1 and HIV etc. (Bradshaw & Venkatesan, 2016, Fragkoudis, Dixon-Ballany et al., 2018, Gagnidze, Hajdarovic et al., 2016). Moreover, nearly half of all emerging viruses are neurotropic viruses (Olival & Daszak, 2005), including the Dengue and Zika viruses (Carod-Artal, 2016, Meyding-Lamade & Craemer, 2018). RABV is a typical neurotropic virus and is the causative agent of rabies disease, a globally well-known and often lethal encephalitis. Therefore, it is urgent to develop new approaches for therapies as well as for cheaper and more effective vaccines against rabies (Fisher & Schnell, 2018, Schnell, McGettigan et al., 2010).

Long non-coding RNAs (lncRNAs) are involved in the development, plasticity, and pathology of the nervous system (Batista & Chang, 2013, Briggs, Wolvetang et al., 2015, Fatica & Bozzoni, 2014, Sun, Yang et al., 2017). Notably, around 40% of lncRNAs detected to date are expressed specifically in the brain (Liu, Wang et al., 2017). Genome-wide association studies (GWASs) and functional studies have associated lncRNAs with neurological diseases including autism spectrum disorders (ASD), schizophrenia, Alzheimer’s disease, and neuropathic pain, among others (Briggs et al., 2015). Mechanistically, it has been shown that lncRNAs can regulate chromatin modifications and gene expression, at both the transcriptional and the post-transcriptional levels (Bonasio & Shiekhattar, 2014, Mercer, Dinger et al., 2009, Wang & Chang, 2011). LncRNAs have recently been shown to regulate innate immune responses by either promoting or inhibiting viral genome replication, highlighting them as a class of novel targets for developing antiviral therapies (Carpenter & Fitzgerald, 2018, Fortes & Morris, 2016, Imamura, Imamachi et al., 2014, Kambara, Niazi et al., 2014, Ma, Han et al., 2017, Ouyang, Hu et al., 2016, Ouyang, Zhu et al., 2014). It is conceivable that antiviral lncRNAs targeting none-innate immune response pathway may exist in neuron cells and brains, which has not been documented yet.

Polycomb repressive complex 2 (PRC2) is a protein complex with an epigenetic regulator function in maintaining the histone modifications that mark transcriptional repression states which are established during early developmental stages (Ringrose, 2017). Some lncRNAs are known to interact with and direct PRC2 towards the chromatin sites of action, thusly defining a trans-acting lncRNA mechanism (Jin, Lv et al., 2018, Rinn, Kertesz et al., 2007). The EZH2 methyltransferase enzyme is the catalytic component of PRC2: it binds RNAs and catalyzes di- or tri-methylation of histone H3 lysine 27 (H3K27me2/3), a modification which leads to the formation of facultative heterochromatin and thus to transcriptional repression (Justin, Zhang et al., 2016, Kasinath, Faini et al., 2018, Margueron & Reinberg, 2011). Many cancers are known to feature very high EZH2 expression levels, so this protein has emerged as an anticancer target for which multiple chemical inhibitors have been developed (Kim & Roberts, 2016, Lee, Yu et al., 2018). It has also been recently reported that inhibitors of the histone methyltransferase activity of EZH2 can suppress infection by several viruses, suggesting a function of EZH2 and/or PRC2 in regulating viral infection (Arbuckle, Gardina et al., 2017). However, it is unclear how this regulation occurs. In general, PRC2 (EZH2) binds different classes of RNAs in a promiscuous manner *in vitro* and in cells, and some lncRNAs such as RepA RNA show *in vitro* specificity with PRC2 (Davidovich, Wang et al., 2015a, Davidovich, Zheng et al., 2013). The specificity of PRC2 (EZH2) interaction with lncRNAs is expected for at least some of its regulation and biological function in living cells, which require further studies (Davidovich, Wang et al., 2015b).

Biochemical studies have established that post-translational modifications (PTM) of EZH2, including phosphorylation and *O*-GlcNAcylation, can regulate its stability (Chu, Lo et al., 2014, Lo, Shie et al., 2018, Wu & Zhang, 2011). NIMA-related kinase (NEK2) was recently shown to phosphorylate EZH2, which protects EZH2 from ubiquitin-dependent proteasome degradation, thereby promoting glioblastoma growth and radio-resistance (Wang, Cheng et al., 2017). LncRNAs have been shown to regulate the stability of proteins such as ZMYND8 and CARM1, expanding the scope of their known regulatory functions (Jin, Xu et al., 2019, Qin, Xu et al., 2019). It was recently reported that a newly identified lncRNA (ANCR) increases the phosphorylation-mediated stability of EZH2 by promoting its interaction with the well-known kinase CDK1 (Li, Hou et al., 2017). However, it remains unclear how lncRNA interacts with proteins to regulate their stability.

Here, we report our discovery of a novel virus-inducible lncRNA (EZH2 degradation-associated lncRNA, EDAL) that we identified via deep RNA-seq of RABV-infected Neuro-2a (N2a) cells. EDAL can inhibit the replication of multiple neurotropic viruses in neuronal cells, including two negative strand RNA viruses-RABV and VSV, a positive strand RNA virus-SFV and a DNA virus-HSV-1, as well as RABV infection in mice. We found that increased EDAL levels reduce the cellular level of EZH2 and of its enzymatic product H3K27me3 epigenetic marks. Mutational analysis, structural prediction, and molecular simulations revealed that a 56-nt functional substructure of EDAL, wherein a helical-loop intimately contacts EZH2 T309 and the surrounding regions. This protein-lncRNA interaction prevents T309 from receiving a previously demonstrated *O*-GlcNAcylation PTM that is known to increase EZH2’s cellular stability. We further show that *Pcp4l1* is a EDAL-regulated gene which encodes a small peptide suppressing RABV, VSV, SFV and HSV-1 infection. Thus, our study reveals a previously unknown lncRNA-PTM-mediated link between host antiviral responses and epigenetic regulation.

## Results

### Identification of a host lncRNA induced by viral infection

We conducted a time-course RNA-seq analysis of cultured N2a cells that were infected with pathogenic RABV (CVS-B2c strain) or were mock infection treated. Subsequently, after a conventional data analysis for differentially expressed mRNA transcripts and a correlation-based analysis to identify time-dependent patterns of transcriptome-wide gene expression changes in response to RABV infection (Appendix Fig S1), we used TopHat2 and Cufflinks (Trapnell, Roberts et al., 2012) to perform a novel lncRNA species prediction, and then conducted a similar differential expression analysis to identify lncRNAs which exhibited significant changes in their accumulation upon RABV infection. This identified 1,434 differentially expressed lncRNAs (Fig 1A). qPCR analysis successfully confirmed the significantly up-regulated expression of ten of the most highly up-regulated of these lncRNAs in response to RABV infection (Fig 1B).

**Figure 1.**
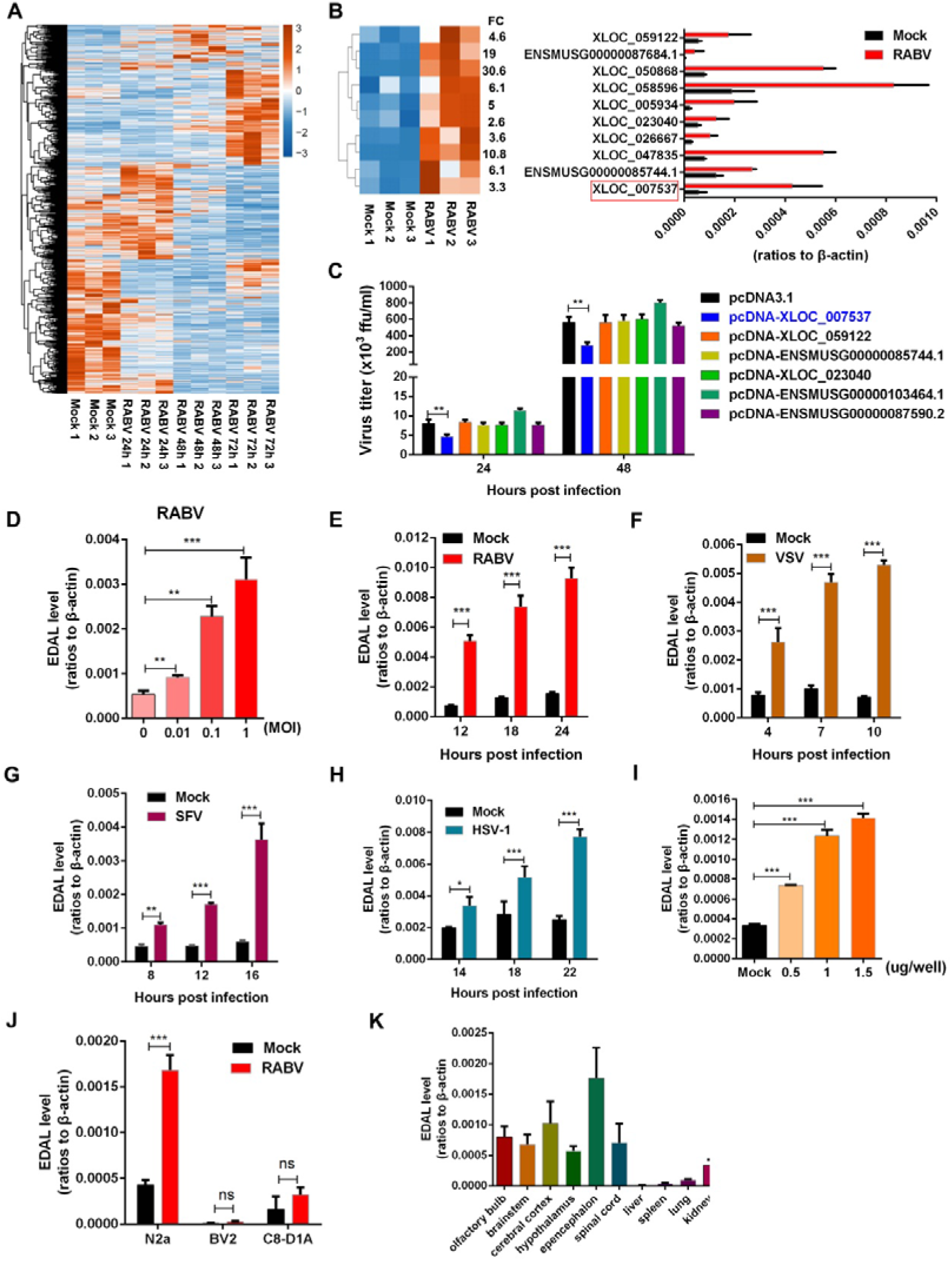
LncRNA EDAL is up-regulated after viral infection. **A.** Total 1434 differentially expressed lncRNAs was identified by RNA-seq analysis in RABV-infected N2a cells compared with mock-infected cells (*n*=3; 2 fold change (FC) and 0.01 *p-value*). These lncRNAs were clustered and shown by heatmap. **B.** Ten of the differentially expressed lncRNAs were selected and clustered in a heatmap (left), the corresponding express level were confirmed by qPCR (right). **C.** The indicated up-regulated lncRNAs were selected and expressed in N2a cells. At 12 h post transfection, the cells were infected with RABV at MOI 0.01 and virus titers in supernatants were measured at indicated time point. **D.** N2a cells were infected with RABV at different MOIs for 24 h and EDAL level was analyzed by qPCR. **E,F,G,H.** N2a cells were infected with RABV (**E**), VSV (**F**), SFV (**G**) or HSV-1 (**H**) at MOI 1 and at indicated time points post infection. EDAL level were determined by qPCR. **I.** N2a cells were transfected with RABV genomic RNA at different doses for 24 h and EDAL level was analyzed by qPCR. **J** The basal or induced level of EDAL (infected with RABV at MOI 1 for 24 h) in different cell lines were determined by qPCR. **K**. The basal level of EDAL in different tissues was analyzed by qPCR. Statistical analysis of grouped comparisons was carried out by student’s t test(\**P* < 0.05; **P<0.01; ***P<0.001). Bar graph represents means ± SD, *n* = 3.

Pursuing the idea that lncRNAs accumulated in response to viral infection may somehow participate in cellular responses to RABV, we cloned six of the strongly up-regulated lncRNAs and overexpressed them in N2a cells; these cells were then infected with pathogenic RABV at a low multiplicity of infection (MOI of 0.01). Excitingly, one of these—XLOC_007537, was predicted to be 1,564 nt in length and to be transcribed from an intergenic locus on chromosome 11—was found to inhibit RABV infection in N2a cells (Fig 1C and Fig EV1A). The 5’ and 3’ boundaries of this XLOC_007537 lncRNA were confirmed by 5’- and 3’-RACE experiments (Fig EV2B). This long intergenic non-coding RNA had no obvious annotation hits after examining its sequence using tools available with the NONCODEv5 (Fang, Zhang et al., 2017), lncRNAdb 2.0 (Quek, Thomson et al., 2015), or LNCipedia 5.0 (Volders, Verheggen et al., 2015) databases. Our PhyloCSF analysis (Lin, Jungreis et al., 2011) yielded a score of -498.50 for this candidate lncRNA (Fig EV1C), strongly reinforcing its non-coding characteristics. Since XLOC_007537 was found to cause EZH2 degradation in the following study, we named it as EZH2 degradation-associated lncRNA (EDAL). EDAL is partially conserved among rats, humans, rhesus, and chimps (Fig EV1D). While RNA fluorescence *in situ* hybridization (FISH) analysis of N2a cells revealed that EDAL occurs in both the cytoplasm and the nucleus, the EDAL signal was stronger in the cytoplasm (Fig EV1E).

### Neuronal cell specific accumulation of EDAL induced by viral infection

We next conducted experiments wherein N2a cells were infected with RABV at different doses for different periods, and EDAL levels were measured via qPCR over a time course of infection. We found that the extent of EDAL up-regulation was dependent on the MOI used for viral infection (Fig 1D), as well as on the infection duration (Fig 1E): increased MOI and increased duration resulted in an increased extent of up-regulation. Bessides RABV, we found several other neutropic viruses, including another negative strand RNA viruses-VSV (Fig 1F and Fig EV2A), a positive strand RNA virus-SFV (Fig 1G and Fig EV2B) and a DNA virus-HSV-1 (Fig 1H and Fig EV2C), could also induce up-regulation of EDAL in a dose- and time-dependent manner. Additional experiments showed that only RABV viral genomic RNA could induce EDAL accumulation: viral proteins, double-stranded RNA (dsRNA), and interferons did not significantly induce EDAL (Fig 1I and Fig EV2D-G).

We then used qPCR to investigate both the basal level and the RABV-induced levels of EDAL in three mouse neuronal cell lines. These experiments revealed that the basal level of EDAL was much higher in N2a cells (neuron cell line) than that in glia cells, including BV2 (microglia cell line) and C8-D1A (astrocyte cell line) cells (Fig 1J). After RABV infection, the level of EDAL in N2a was significantly up-regulated, while no significant change in the EDAL level was detected in BV2 or C8-D1A cells (Fig 1J). Furthermore, EDAL levels were much higher in brains and spinal cords than in the spleen, liver, or lung (Fig 1K).

### EDAL inhibits viral replication

We next transfected N2a cells with pcDNA3.1 plasmid expressing either EDAL (pcDNA-EDAL) or an EDAL-specific small interfering RNA (siEDAL) and then verified that EDAL was appropriately expressed or specifically silenced in N2a cells (Fig EV3A and 3B). We also confirmed that overexpression or silencing of EDAL did not affect cell viability (Fig EV3C and 3D). Next, we transfected N2a cells with the EDAL expression plasmid and then infected them with RABV at 12 hours (h) post transfection. The viral titer in the supernatant of RABV-infected cells transfected with the pcDNA-EDAL vector was 8-fold lower than the titer of control cells transfected with the empty vector pcDNA3.1 at 48 h post infection (hpi). At 72 hpi, the same trend was apparent, but the difference was 4.5-fold (Fig 2A).

**Figure 2.**
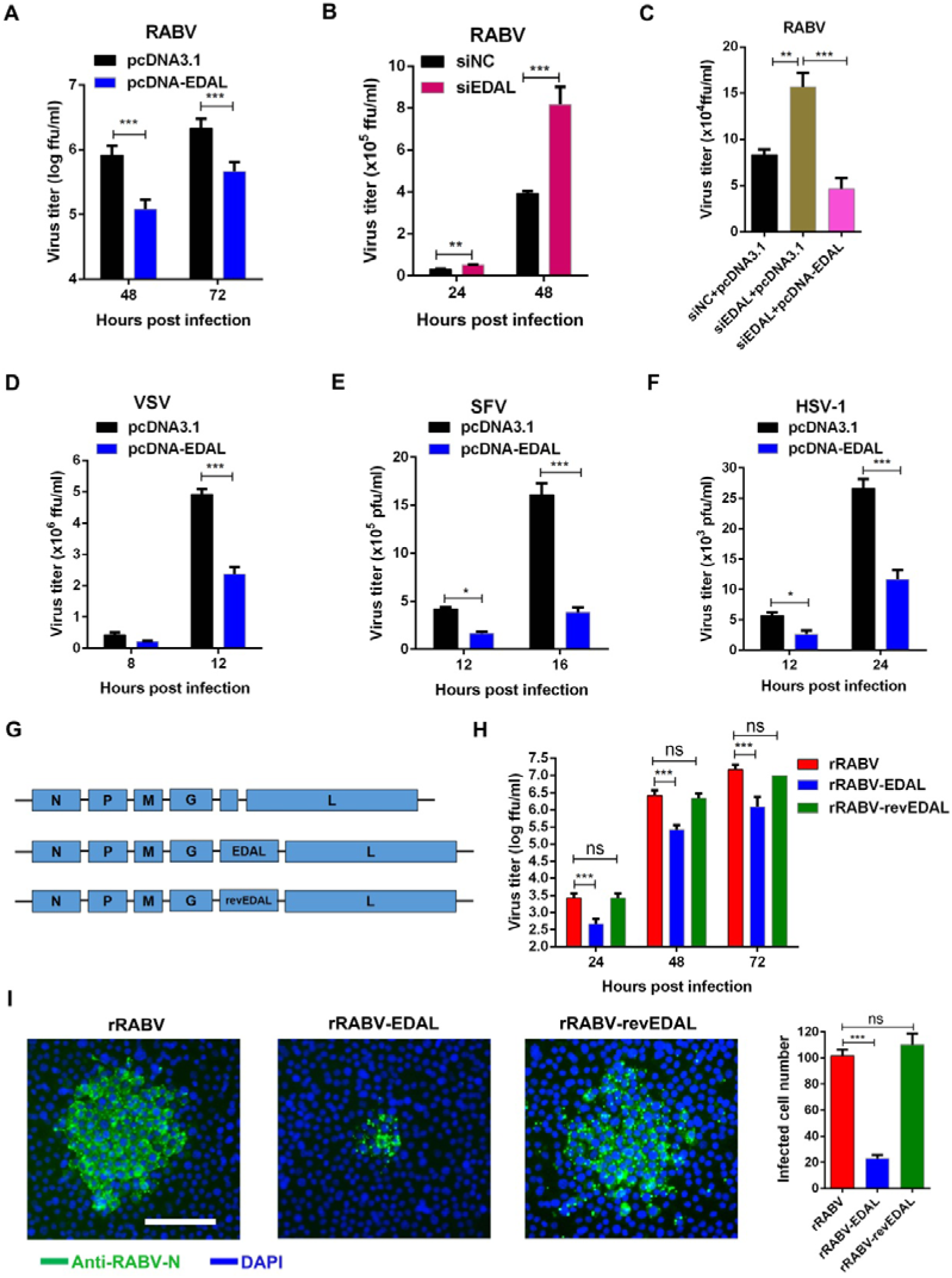
EDAL inhibits viral replication in neuronal cells. **A.** N2a cells were transfected with pcDNA3.1 or pcDNA-EDAL, then at 12 h post transfection the cells were infected with RABV at MOI 0.01 and virus titers were measured at indicated time points. **B.** N2a cells were transfected with EDAL specific siRNA (siEDAL) and at 12 h post transfection, the cells were infected with RABV at MOI 0.01 and virus titers were measured at indicated time points. **C.** N2a cells were transfected with siEDAL or siNC (negative control) for 8 h and then transfected with pcDNA3.1 or pcDNA-EDAL. At 12 h post transfection, the cells were infected with RABV at MOI 0.01 for 24 h and virus titers in the cell supernatant were measured. **D.** N2a cells were transfected with pcDNA3.1 or pcDNA-EDAL, then at 12 h post transfection the cells were infected with VSV at MOI 0.01 and virus titers were measured at indicated time points. **E.** N2a cells were transfected with pcDNA3.1 or pcDNA-EDAL, then at 24 h post transfection the cells were infected with SFV at MOI 0.01 and virus titers were measured at indicated time points. **F.** N2a cells were transfected with pcDNA3.1 or pcDNA-EDAL, then at 12 h post transfection the cells were infected with HSV-1 at MOI 0.01 and virus titers were measured at indicated time points. **G,H.** EDAL and reverse EDAL (revEDAL) were inserted into the genome of a recombinant RABV (rRABV), named rRABV-EDAL and rRABV-revEDAL respectively (**G**), and their growth kinetics in N2a cells (MOI=0.01) were compared (**H**). **I.** N2a cells were infected with rRABV, rRABV-EDAL or rRABV-revEDAL at MOI 0.005 for 48 h and the viral spread were compared by calculating the cell numbers within the fluorescence focus. Scale bar, 50 μm. Statistical analysis of grouped comparisons was carried out by student’s t test(\**P* < 0.05;**P<0.01; ***P<0.001). Bar graph represents means ± SD, *n* = 3.

The impact of EDAL silencing on virus titer was assessed using direct immunofluorescence assays with an antibody against the RABV N protein, which allowed calculation of the number of living RABV particles according to the number of immunofluorescent foci (Tian, Luo et al., 2016). Excitingly, and consistent with a virus-replication-inhibiting function for EDAL in N2a cells, when the expression of EDAL was silenced by siEDAL, the RABV titer increased by around 2-fold compared to the siRNA control cells at 48 hpi. (Fig 2B), and the impact of siEDAL silencing was removed by subsequent overexpression of EDAL (Fig 2C). Interestingly, A similar trend of reduced viral titers in cells transfected with the EDAL plasmid was observed in VSV, SFV and HSV-1-infected cells (Fig 2D-F).

To further explore a role for EDAL in somehow inhibiting viral replication, we next developed a series of recombinant viruses for later experiments with live mice. Specifically, we here used a recombinant RABV (rRABV) virus that was derived from the CVS-B2c strain, and used three different viral constructs: unaltered rRABV, rRABV harboring the EDAL sequence (rRABV-EDAL), and rRABV harboring the reverse complement sequence of EDAL (rRABV-revEDAL) (Fig 2G). Virus growth kinetics experiments with N2a cells showed that the virus titer was significantly lower in the rRABV-EDAL infected cells than both the rRABV-infected cells and the rRABV-revEDAL-infected cells (Fig 2H).

We also analyzed the capacity of the recombinant viruses to spread between infected cells and neighboring cells, the infected N2a cells were covered by low melting agar to inhibit the virus release into the supernatant (Tian et al., 2016). The rRABV-EDAL recombinant virus yielded much smaller fluorescent foci than rRABV and rRABV-revEDAL in the neighboring N2a cells (Fig 2I, left) at 48 hpi, and the fluorescent foci we observed in the rRABV-EDAL-infected samples comprised significantly fewer cells than the fluorescent foci present in the rRABV or rRABV-revEDAL samples (Fig. 2I, right).

### EDAL reduces RABV pathogenicity in vivo

To investigate the role of EDAL in RABV infection *in vivo*, we compared the pathogenicity of rRABV, rRABV-EDAL, and rRABV-revEDAL in the C57BL/6 mouse model. Mice were infected intra-nasally (i.n.) with rRABV, rRABV-EDAL, or rRABV-revEDAL (100 FFU). The mice infected with rRABV and rRABV-revEDAL exhibited decreased body weights starting from 7 to 9 days post infection (dpi), and these decreases became significant between 9 and 14 dpi. In contrast, the body weight of mice infected with rRABV-EDAL only exhibited a slight decrease between 10-14 dpi (Fig 3A). Moreover, the rabies symptoms (including weight loss, ruffled fur, body trembling, and paralysis) of the symptomatic rRABV- and rRABV-revEDAL-infected mice appeared at 7 dpi, and became exacerbated until death at 14 dpi, whereas symptomatic mice infected with rRABV-EDAL had only mild symptoms which occurred from 9 to 15 dpi (Fig 3B). Among all mice, 70% of the mice infected with rRABV-EDAL survived, compared with only 20% and 10% survival ratio for rRABV- and rRABV-revEDAL-infected mice, respectively (Fig 3C).

**Figure 3.**
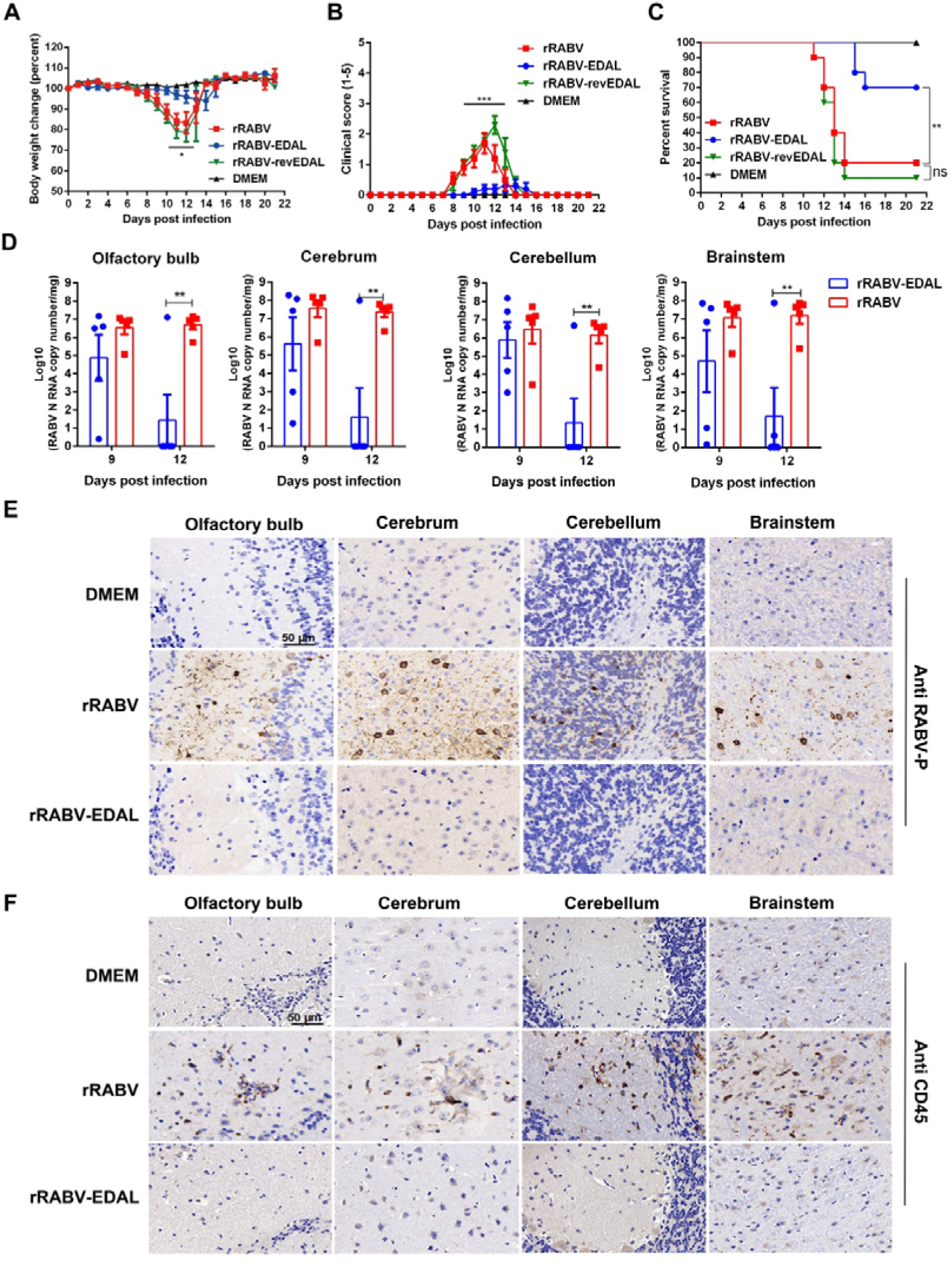
EDAL attenuates RABV pathogenicity *in vivo*. **A,B,C.** Female C57BL/6 mice (8-week-old, n=10) were infected intranasally with 100 FFU rRABV, rRABV-EDAL, or rRABV-reEDAL, or mock infected. Body weight change (**A**), clinical score (**B**) and survival ratio (**C**) were monitored daily for continuous 3 weeks. (means ± SEM; **P<0.01; body weight change and clinical score was analyzed by Two-way ANOVA test; survival ratio was analyzed by log rank test). **D.** At indicated time points, the brains from the infected mice were collected for analyzing the level of RABV N mRNA by qPCR. (n=5; means ± SEM; **P<0.01 by student’s Two-way ANOVA test). **E,F.** At 12 dpi, the brains were collected, resolved by paraffin sections, and analyzed by immunohistochemistry by staining with antibodies against RABV P (**E**) or CD45 (**F**). Scale bar, 50 μm.

To quantify the viral load in rRABV and rRABV-EDAL infected brains, the RABV *N* mRNA level in different encephalic regions was analyzed by qPCR after i.n. infection with 100 FFU of different viruses. At 12 dpi, we observed dramatically reduced RABV *N* mRNA levels in rRABV-EDAL-infected vs. rRABV-infected mice: specifically, these reductions were observed in the olfactory bulb, cerebrum, cerebellum, and brain stem regions (Fig 3D). Further immunohistochemistry analysis of the RABV P protein (Fig 3E) and CD45-positive cells (Fig 3F) in various brain regions showed that, unlike rRABV-infected brains, almost no viral antigen or virus-induced inflammation could be observed in rRABV-EDAL-infected mouse brains at 12 dpi. Collectively, these results establish that EDAL can dramatically inhibit intranasal-inoculation-induced RABV infection in mice.

### EDAL decreases H3K27me3 levels by promoting lysosome-mediated EZH2 degradation

Having demonstrated that RABV infection induces the accumulation of EDAL and established that EDAL can restrict RABV replication *in vitro* and *in vivo*, we were interested in potential mechanism(s) through which EDAL may exert its antiviral effects. We have for some time been interested in the potential contributions of epigenetic regulation on host responses to neurotropic viruses, and we noted that the N2a cells transfected with the pcDNA3.1 plasmid expressing pcDNA-EDAL had significantly decreased levels of histone methylation. Specifically, immunoblotting experiments with an antibody against the H3K27me3 tri-methylation mark revealed that cells with the empty control plasmid had a signal for this histone methylation of the N-terminal tail of the core histone H3 that was 1.35 times as strong as the signal for cells with the pcDNA-EDAL plasmid (Fig 4A).

**Figure 4.**
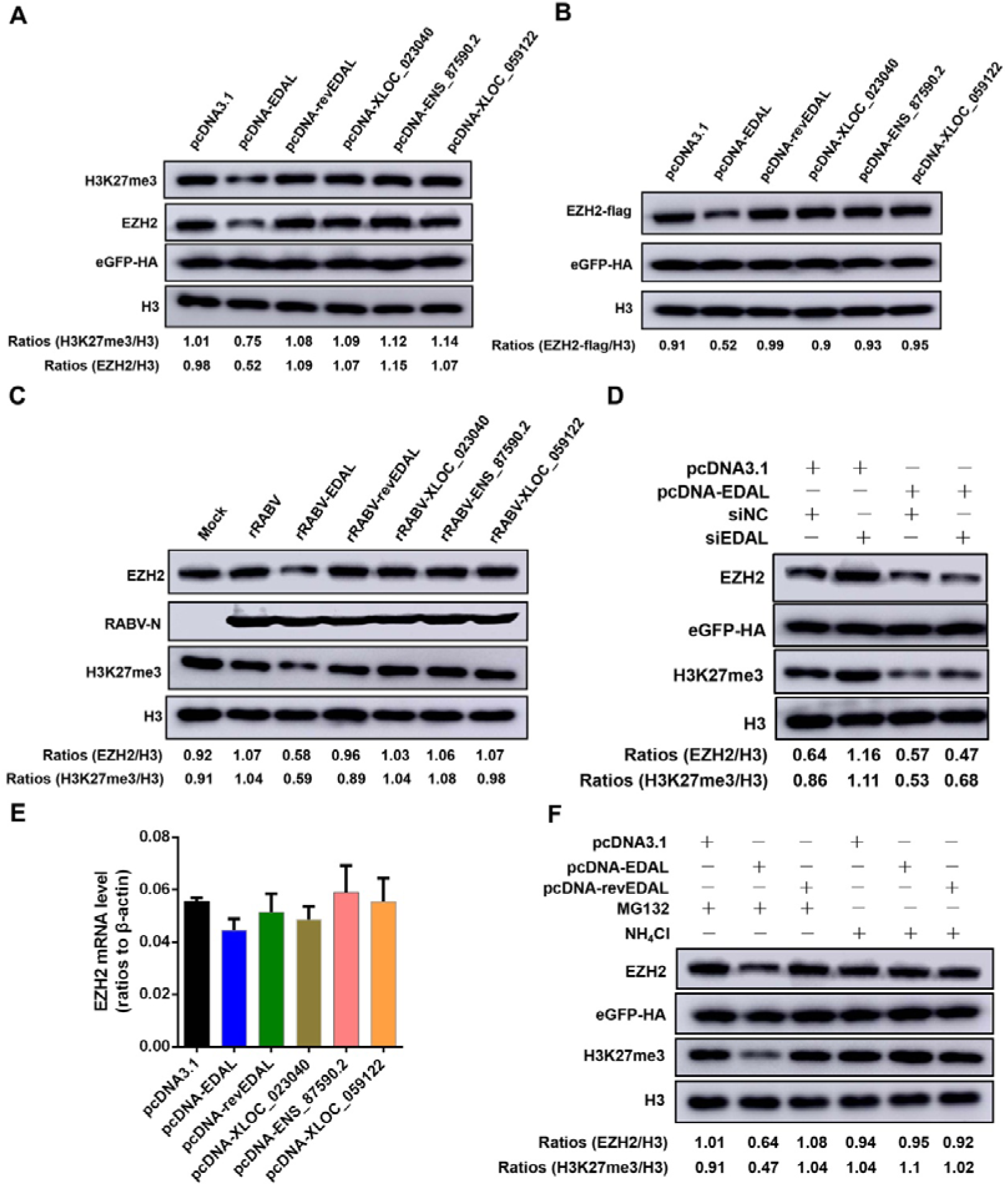
EDAL down-regulates H3K27me3 level by causing the degradation of EZH2. **A.** EDAL, reverse EDAL (revEDAL), XLOC_023040, ENSMUSG00000087590.2 (ENS_87590.2) or XLOC_059122 was overexpressed in N2a cells for 48 h and then EZH2 or H3K27me3 level were resolved by Western blotting. The plasmid pCAGGS-eGFP containing a HA tag was used as a transfection control. **B.** N2a cells were transfected with pcDNA3.1, pcDNA-EDAL, pcDNA-revEDAL, pcNDA-XLOC_023040, pcDNA-ENS_87590.2, or pcDNA-XLOC_059122, and pCAGGS-EZH2-FLAG and pCAGGS-eGFP-HA (transfection control). EZH2-FLAG levels were measured by Western blotting and normalized to H3. **C.** N2a cells were infected with rRABV, rRABV-EDAL, rRABV-revEDAL, rRABV-XLOC_023040, rRABV-ENS_87590.2 or rRABV-XLOC_059122 at MOI 3. At 36 hpi, the EZH2 and H3K27me3 level was resolved by Western blotting and normalized to H3. **D.** N2a cells were transfected with siEDAL or siNC (negative control) for 8 h and then transfected with pcDNA3.1 or pcDNA-EDAL. Then EZH2 and H3K27me3 level was resolved by Western blotting and normalized to H3. **E.** N2a cells were transfected with pcDNA3.1, pcDNA-EDAL, pcDNA-revEDAL, pcNDA-XLOC_023040, pcDNA-ENS_87590.2, or pcDNA-XLOC_059122. The mRNA levels of EZH2 were analyzed by qPCR. (n=3). **F.** pcDNA3.1, pcDNA-EDAL or pcDNA-revEDAL was transfected into N2a cells. The specific inhibitors for proteasome and lysosome, MG132 (10 µM) and NH_4_Cl (5 mM), were applied. Then EZH2 and H3K27me3 level was analyzed by Western blotting and normalized to H3.

To confirm an impact specifically from EDAL on the observed reduction in the H3K27me3 tri-methylation level, we evaluated three other lncRNAs from our dataset which were induced by RABV, namely XLOC_023040, ENSMUSG00000087590.2 (ENS_87590.2), and XLOC_059122 mentioned in Fig. 1C. Notably, the expression of these lncRNAs did not change the H3K27me3 tri-methylation level (Fig 4A), strongly supporting the specificity of EDAL in exerting this inhibitory effect. These results led us to speculate that EDAL may interfere with viral replication via alteration of histone methylation.

It is now understood that PRC2 mediates the H3K27me3 tri-methylation process (Simon & Kingston, 2009), so we performed additional immunoblotting with an antibody against EZH2—the enzymatic subunit of PRC2 responsible for its methyl-transferase activity. As with the signal for H3K27me3 tri-methylation, we observed weaker signals for EZH2 in cells with the plasmid for pcDNA-EDAL compared to controls (Fig 4A and B). We next used the recombinant viruses that we used for mice infection (Fig 3) to repeat the above experiments, and the same decreasing trend was observed in N2a cells infected with the rRABV-EDAL virus (Fig 4C). Moreover, no such decreases in the H3K27me3 tri-methylation signal or the EZH2 protein level were observed upon expression of revEDAL or the three aforementioned lncRNAs (Fig 4C), again highlighting an apparently specific contribution of EDAL to the reduced levels of H3K27me3 and its catalyst EZH2.

To further determine the impact of EDAL on the H3K27me3 tri-methylation signal and/or the EZH2 protein level, N2a cells were transfected with siEDAL. Consistently, silencing of EDAL enhanced the levels of both EZH2 and H3K27me3 in N2a cells (Fig 4D), and overexpression of EDAL counteracted the elevated EZH2 level induced by siEDAL (Fig 4D). Importantly, we also found that the EZH2 protein level, but not the *EZH2* mRNA level, was reduced by EDAL—and noted that expression of revEDAL or other three control lncNRAs did not affect the protein or the mRNA level for EZH2 (Fig 4E)—results clearly suggesting that the impact of EDAL on EZH2 accumulation occurs at the protein level.

We therefore suspected that an EDAL–EZH2 interaction might somehow promote the degradation of EZH2, thereby reducing the overall cellular capacity for its methyltransferase activity, potentially explaining the observed reduction in H3K27me3 tri-methylation. To test this hypothesis, we treated cells with compounds that inhibit the protein degradation functions of proteasomes (MG132) or lysosomes (NH_4_Cl), and then assayed the EZH2 protein accumulation and the H3K27me3 tri-methylation level upon EDAL expression. These experiments showed that NH_4_Cl but not MG132 treatment restored the EZH2 protein and H3K27me3 tri-methylation levels, results supporting that EDAL somehow causes EZH2 degradation via the lysosomal degradation pathway (Fig 4F).

### A 56 nt 5’ segment is responsible for EDAL’s antiviral activity

Although not necessarily conserved, secondary structures are thus far good candidates for identification of functional elements of lncRNAs (Bonasio & Shiekhattar, 2014, Johnsson, Lipovich et al., 2014, Mercer & Mattick, 2013, Rivas, Clements et al., 2017). Seeking to identify secondary structures of EDAL that affect its specific interaction with EZH2, predictions using the RNAstructure 5.3 program indicated that EDAL could be divided into four major sub-structures, each containing a number of base-paired structures and hairpin structures (Fig 5A). We cloned the segments corresponding to the four sub-structures (EDAL-1, EDAL-2, etc.) into pcDNA3.1, and then each of the four segments was individually expressed in N2a cells, followed by immunoblotting-based evaluation of the EZH2 protein and H3K27me3 tri-methylation levels. Interestingly, the first truncated segment (EDAL-1) located at the 5’ end of EDAL, but none of the other three segments, significantly reduced both the EZH2 and H3K27me3 levels (Fig 5B). Consistent with a specific impact from this EDAL sub-structure, only EDAL-1 restricted RABV replication in N2a cells (Fig 5C).

**Figure 5.**
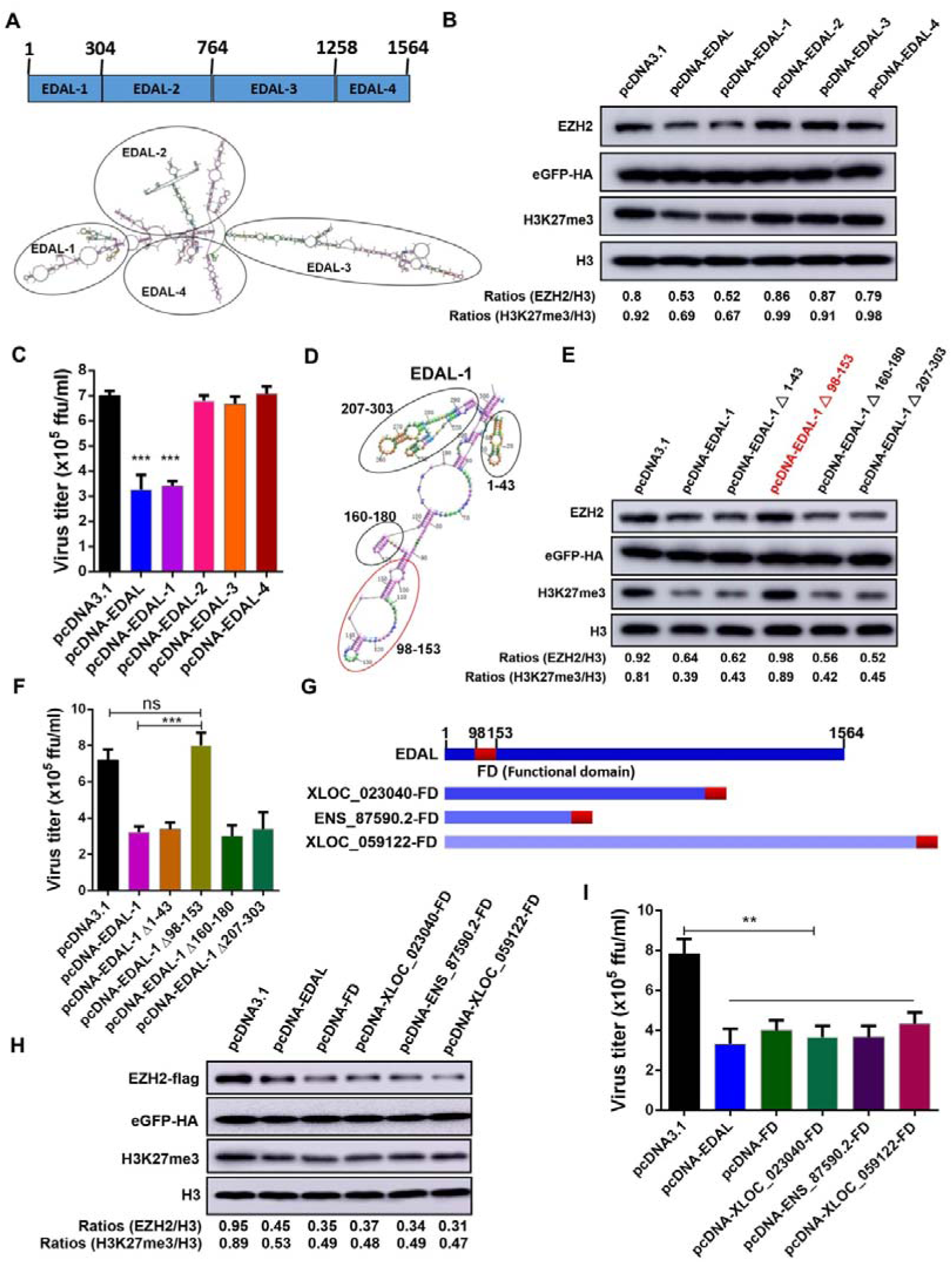
The 56-nt portion of EDAL in 5’ end carries the antiviral function. **A.** EDAL secondary structure was predicted by RNAstructure Version 5.8 software (http://rna.urmc.rochester.edu/rnastructure.html). EDAL was divided into four sections based on sub-structures: EDAL-1(1-304 nt), EDAL-2 (305-764 nt), EDAL-3 (765-1258 nt) and EDAL-4 (1259-1564 nt). **B.** The full-length EDAL and its truncations were separately transfected into N2a cells for 48 h. The EZH2 and H3K27me3 level was resolved by Western blotting and the ratio normalized to H3 was calculated. **C.** The full-length EDAL and its truncations were expressed in N2a cells for 12 h and then the cells were infected with RABV at MOI 0.01. At 48 hpi, the virus titers in the cell supernatant were measured. **D,E.** Four sections within EDAL-1 were selected based on the secondary structures (**D**). The four truncations EDAL-1 deleting 1-43 nt (EDAL-1 1-43), 98-153 nt (EDAL-1 98-153), 160-180 nt (EDAL-1 160-180) and 207-303 nt (EDAL-1 207-303) were cloned into pcDNA3.1, respectively. The different truncations as well as full length EDAL-1 were overexpressed in N2a cells for 48 h. Then EZH2 and H3K27me3 level was resolved by Western blotting and normalized to H3 (**E**). **F.** N2a cells were transfected with pcDNA3.1, pcDNA-EDAL-1 or different truncations of EDAL-1 for 12 h. Then the cells were infected with RABV at MOI 0.01 and the virus titers in supernatant were measured at 48 hpi. **G,H.** The functional domain (FD) of the 56-nt portion of EDAL was cloned into pcDNA3.1 or fused with 3’ end of the other three control lncRNAs (**G**). Then these lncRNAs were transfected together with pCAGGS-EZH2-flag into N2a cells for 48 h. EZH2 and H3K27me3 level were analyzed by Western blotting and normalized to H3 (**H**). **I.** N2a cells were transfected with pcDNA3.1, pcDNA-EDAL or different recombinant lncRNAs for 12 h. Then the cells were infected with RABV at MOI 0.01 and the virus titers in supernatant were measured at 48 hpi. Statistical analysis of grouped comparisons was carried out by student’s t test(**P<0.01; ***P<0.001). Bar graph represents means ± SD, *n* = 3.

To pinpoint the specific fragment capable of exerting the antiviral function, EDAL-1 was assessed as four separate truncation segments (EDAL-1 △ 1-43, EDAL-1 △ 98-153, EDAL-1 △ 160-180 and EDAL-1 △ 207-303) (prepared as depicted in Fig 5D). Each of the EDAL-1 variants were assessed in N2a cells: only EDAL-1 △ 98-153 failed to decrease the EZH2 and H3K27me3 levels and failed to inhibit rRABV replication (Fig 5E and F).

To confirm that EDAL 98-153 nt can inhibit RABV infection, this 56 nt segment was expressed by itself and as a fusion with the 3’ end of the three aforementioned lncRNAs (i.e., from our experiments to successfully demonstrate the specificity of EDAL’s antiviral effects) (Fig 5G). As expected, the fragment alone and the three fusion lncRNAs reduced the EZH2 and H3K27me3 levels (Fig 5H) and also reduced RABV replication (Fig 5I). These results establish that the 56 nt segment at the 98-153 position of the 5’ end of EDAL is essential for the EZH2-mediated antiviral effects we observed in neuronal cells.

### EDAL reduces EZH2 stability by impeding an O-GlcNAcylation PTM at the T309 site

Previous studies have revealed that phosphorylation and *O*-GlcNAcylation can influence the stability of EZH2 (Chu et al., 2014, Lo et al., 2018, Wu & Zhang, 2011). At least two phosphorylation sites among human EZH2, T345 and T487, were shown to affect its stability (Wu & Zhang, 2011). However, we found that EDAL could still cause the degradation of murine EZH2 when the corresponding phosphorylation sites were mutated to T341A and T485A, (Fig EV4A), indicating that EDAL does not apparently impair the phosphorylation of EZH2.

There are five known *O*-GlcNAcylation sites (S73, S76, S84, T313, and S729) in human EZH2 that can regulate EZH2 stability and enzymatic activity (Chu et al., 2014, Lo et al., 2018). Based on the sequence alignment between human and murine EZH2, we found that S73, S75, T309, and S725 are potential *O*-GlcNAcylation sites of murine EZH2 (Fig EV4B). We mutated each of the potential *O*-GlcNAcylation sites of murine EZH2 and then co-transfected these mutant variants together with pcDNA3.1, pcDNA-EDAL, or pcDNA-revEDAL in N2a cells. We found only T309A mutation lost the EDAL-promoted EZH2 degradation (Fig 6A), while there was no significant difference in the extent of degradation among the wild type, S73A, S75A, or S725A variants of EZH2 (Fig 6A). We observed the same trends for EZH2 variants bearing multiple mutations: a S73/S75/S725 triple-alanine-mutant did not affect EDAL-promoted EZH2 degradation, whereas EDAL lost its impact on the degradation of a tetra-alanine EZH2 variant with mutation of position 309 (Fig 6B). These results together indicated that EDAL mediated EZH2 degradation via specifically blocking T309 *O*-GlcNAcylation site.

**Figure 6.**
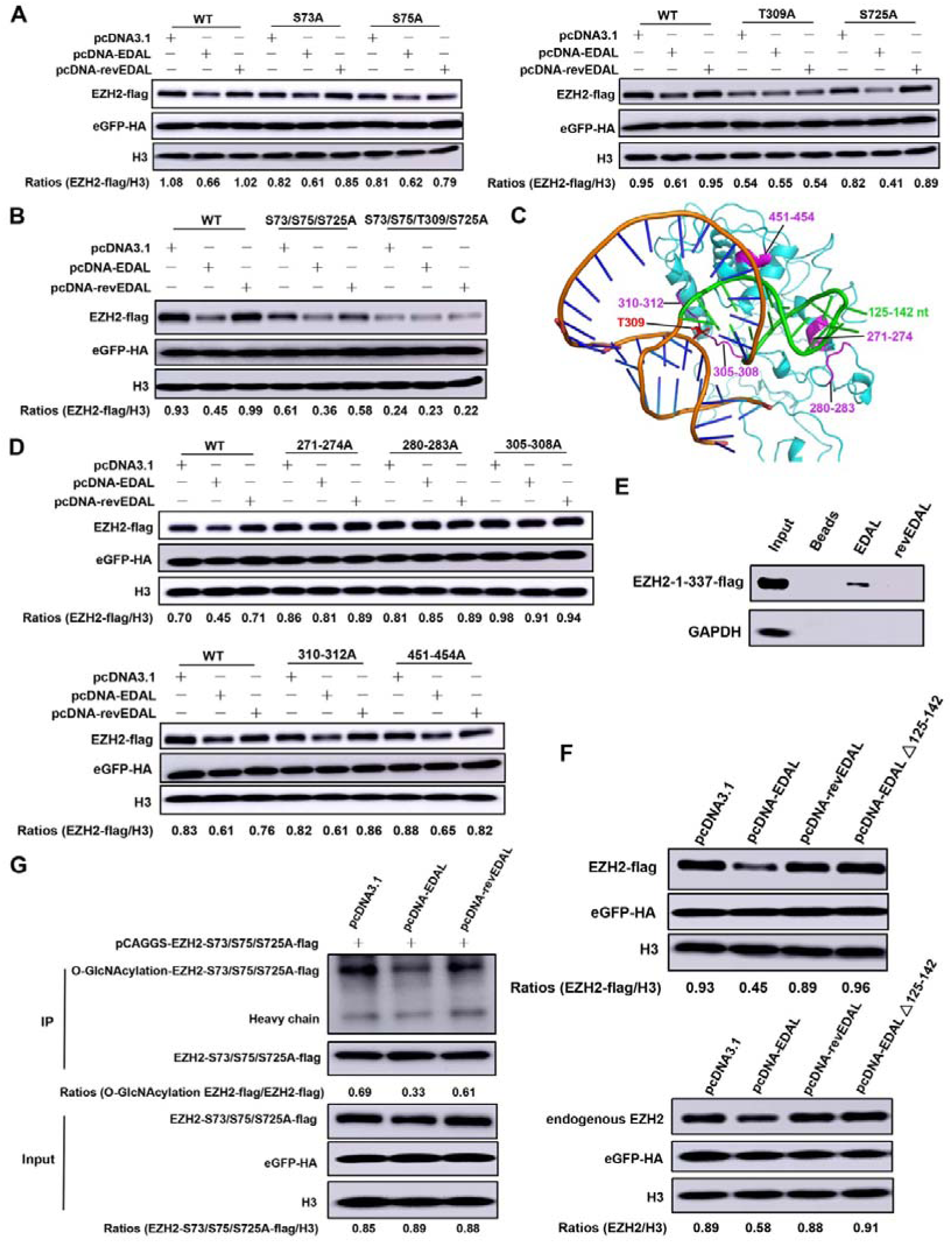
EDAL promotes EZH2 degradation via impeding the *O*-GlcNAcylation at T309 site. **A.** The potential *O*-GlcNAcylation sites of murine EZH2 was individually mutated and expressed together with EDAL or revEDAL in N2a cells for 48 h. Then EZH2 level was analyzed by Western blotting and normalized to H3. **B.** The potential *O*-GlcNAcylation sites of murine EZH2 was mutated and co-expressed together with EDAL or revEDAL in N2a cells for 48 h. Then EZH2 level was analyzed by Western blotting and normalized to H3. **C.** Murine EZH2 3D structure was predicted with SWISS-MODEL (https://swissmodel.expasy.org/interactive) based on human EZH2 3D structure (PDB code: 5HYN). EDAL-FD 3D structure model was predicted with RNAComposer (http://rnacomposer.ibch.poznan.pl/). The interaction between EDAL functional domain (98-153 nt) and EZH2 was predicted by 3dRPC. The predicted interactional residues among EZH2 were marked with magenta color and among EDAL with green color. **D.** The predicted interaction residues of EZH2 were mutated and cloned into pCAGGS vector, and then co-transfected with pcDNA3.1, pcDNA-EDAL or pcDNA-revEDAL in N2a cells for 48 h. Then EZH2 level was analyzed by Western blotting and normalized to H3. The plasmid pCAGGS-eGFP containing a HA tag was used as a transfection control. **E.** RNA pull-down analysis of the binding of EDAL or revEDAL to EZH2-1-337-flag. **F.** EDAL deleting 125-142 nt (EDAL △ 125-142) was cloned into pcDNA3.1 (pcDNA-EDAL △ 125-142). Then pcDNA3.1, pcDNA-EDAL, pcDNA-revEDAL and pcDNA-EDAL △ 125-142 were individually or together with pCAGGS-EZH2-flag transfected into N2a cells for 48 h. Then the overexpressed EZH2 (EZH2-flag) and endogenous EZH2 level was resolved by Western blotting and normalized to H3. **G.** The plasmid expressing EZH2-S73/S75/S725A-flag was co-transfected with pcDNA3.1, pcDNA-EDAL or pcDNA-revEDAL in N2a cells and treated with NH_4_Cl (5 mM) for 48 h. Then the *O*-GlcNAcylation level of EZH2-S73/S75/S725-flag was analyzed by Western blotting.

In order to further pursue the EDAL-EZH2 interactions which may contribute to the EDAL specific blocking of the EZH2 T309 *O*-GlcNAcylation, we decided to predict the interaction sites between the 56-nt antiviral EDAL substructure and EZH2. RNA tertiary structure prediction revealed a tertiary structure for the 56-nt antiviral RNA segment: the helix-loop tertiary structure folded by the 18-nt terminal hairpin corresponding to 125-142 of EDAL was packed on the second helix folded by the stem base-paired structure, and most of the two structural components were free for contacting other partners (Fig 6C). We then conducted for molecular docking using the 3dRPC program taking the advantage of recently published tertiary structures for EZH2 (Huang, Li et al., 2016, Huang, Li et al., 2018, Justin et al., 2016, Kasinath et al., 2018). Among the top scored structures, one showing that the 18-nt terminal helix-loop tertiary structure was intimately interacted with EZH2 residues at positions 271-274, 280-283, 305-308, 310-312, and 451-454 (Fig 6C). To validate these predicted interactions, we mutated all these EDAL interacting residues in EZH2 to alanine (A). We co-transfected N2a cells with plasmids expressing wild type EZH2 and EZH2 mutant variants together with the pcDNA3.1, pcDNA-EDAL or pcDNA-revEDAL plasmids. The results revealed a striking difference: in the presence of EDAL, there was no obvious reduction in the levels of the EZH2 variants bearing alanine substitution mutations at the 271-274, 280-283, or 305-308 positions, whereas there was obvious degradation of WT EZH2 and the other variants (Fig 6D). Thus, the cellular stability of EZH2 is directly affected by an interaction between EDAL and the EZH2 residues at positions 271-274, 280-283, and 305-308. Previous studies have demonstrated that the binding region between human EZH2 and many reported lncRNAs was the segment of 343-368 aa (Kaneko, Li et al., 2010), and the corresponding region in murine EZH2 was between 338 and 364 aa determined by sequence comparison. However, our results indicate that EDAL binds to murine EZH2 in the region of 271-274, 280-283 and 305-308. In order to verify these binding sites between murine EZH2 and EDAL, we truncated murine EZH2 into 1-337aa and cloned the truncated fragment into pCAGGS vector. Then we confirmed that the specific EDAL interaction sites on EZH2 are in its N-terminal region (1-337 aa) using an RNA pull-down analysis (Fig 6E). Reciprocally, the expression of an EDAL variant lacking the 18-nt terminal hairpin segment (125-142 nt) lost the ability to promote the degradation of both over-expressed and endogenous EZH2 (Fig 6F).

The molecular docking and validation experiments supported a model that EDAL can specifically binds to EZH2 at T309 *O*-GlcNAcylation site. We therefore speculated that EDAL binding might impair the *O*-GlcNAcylation at T309 site, potentially preventing an EZH2-stability-promoting effect associated with this PTM. Pursuing this, we evaluated the effect of EDAL expression on the *O*-GlcNAcylation level of EZH2 at the T309 site. To exclude the impact of other *O*-GlcNAcylation sites on the detected level of EZH2 *O*-GlcNAcylation, pCAGGS-EZH2-S73/S75/S725A-flag plasmid was transfected together with pcDNA3.1, pcDNA-EDAL, or pcDNA-revEDAL into N2a cells, and then the *O*-GlcNAcylation level on the EZH2-S73/S75/S725A-flag fusion protein was measured post treatment with NH_4_Cl. Interestingly, we found that expression of EDAL dramatically reduced the *O*-GlcNAcylation level of EZH2 (Fig 6G). These results support that EDAL specifically contacts T309, shielding T309 from *O*-GlcNAcylation.

### The EZH2 inhibitor gsk126 protects neuronal cells from viral infection

If EDAL’s antiviral effects are indeed mediated by its reduced EZH2 methyltransferase activity, then we could anticipate that chemical inhibition of EZH2 should cause antiviral effects. Gsk126 is a specific inhibitor of EZH2 methyltransferase activity (Mccabe, Ott et al., 2012), and we evaluated the effects of gsk126 on RABV and VSV replication in N2a cells. After testing toxicity (Appendix FigS2A) and identifying a suitable working concentration of gsk126 (Appendix Fig S2B), we pretreated N2a cells with 4 µmol (µM) gsk126 and then infected them with rRABV, or VSV. The replication of both rRABV (Appendix Fig S2C) and VSV (Appendix Fig S2D) was significantly decreased by treatment with gsk126, results which reinforce a specific role for EZH2’s methyltransferase activity on the antiviral effects we observed in N2a cells and which demonstrate proof-of-concept for a therapeutic strategy against a neurotropic virus.

### EDAL restricts viral replication by up-regulation of an antiviral peptide PCP4L1

Next we attempt to identify the genes which might be up-regulated by EDAL via decreasing H3K27me3 levels. N2a cells were transfected with pcDNA-EDAL or pcDNA3.1, and then infected with RABV at MOI 1. At 48 hpi, the poly(A)-RNA was isolated for deep sequencing. A cut-off of 0.05 FDR resulted in a total of 75 up-regulated genes (Fig 7A). We next wanted to identify the direct EDAL targets among genes regulated by EDAL *in trans*. We turned our attention to the altered H3K27me3 modification as an additional selection criterion for EDAL to induce EZH2 degradation and reduce H3K27me3 level. Chromatin Immunoprecipitation Sequencing (ChIP-seq) was performed by using anti-H3K27me3 antibody to profile the distribution of H3K27me3 marks on the genome of N2a cells upon transfection with pcDNA-EDAL or control plasmids, and then the data were summarized in Appendix Table S1. Analysis of H3K27me3 peaks indicative of the epigenetic silencing positions revealed many fewer peaks—11,918 vs. 59,706—in EDAL overexpressed samples compared with the samples transfected with empty control plasmids, consistent with the EDAL-reduced cellular level of H3K27me3. In total, 2026 genes lost H3K27me3 mark and only 167 genes gained after EDAL overexpression (Fig 7B). Most EDAL-upregulated genes naturally did not contain H3K27me3 mark, consistent with a recent report that many H3K27me3 marks in adult mice is not related to transcriptional regulation (Jadhav, Nalapareddy et al., 2016).

**Figure 7.**
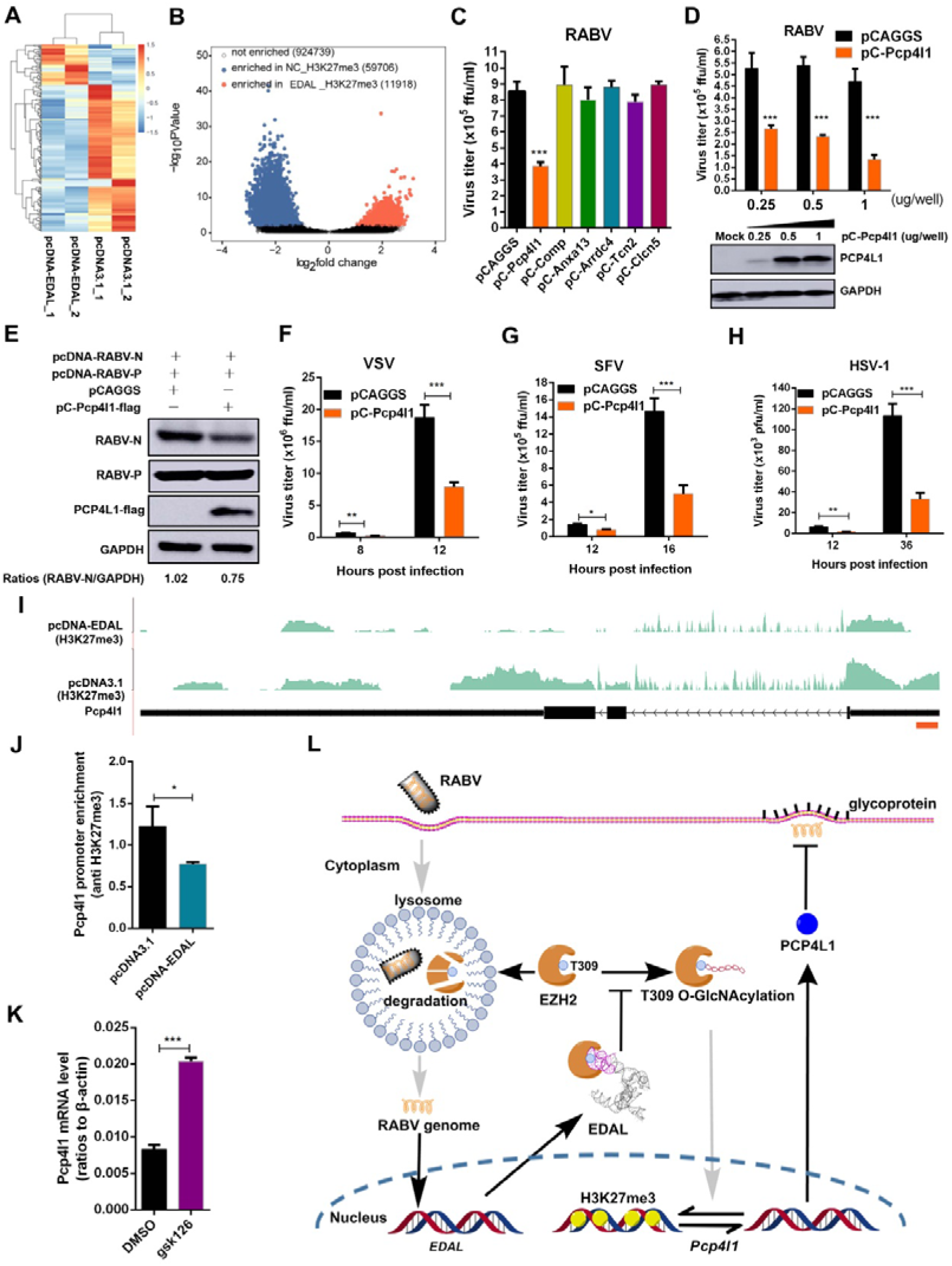
EDAL restricts viral replication by up-regulation of *Pcp4l1*. **A.** N2a cells were transfected with pcDNA3.1 or pcDNA-EDAL for 12 h and then infected with RABV at MOI 1 for 48 h. Total RNA was isolated and subjected to RNA-seq analysis (*n*=2; 2 fold change (FC) and 0.01 *p-value*). **B.** N2a cells were transfected with pcDNA3.1 or pcDNA-EDAL for 48 h and then ChIP-seq analysis was performed. Volcano plot showed the peaks enriched in negative control (NC) cells and EDAL overexpression cells. X axis was the log2 ratio of EDAL versus NC signals for each peak, and Y axis was the significance of the differences (−log10 (*P-values*)). **C.** Six up-regulated and loss of H3K27me3 mark genes were cloned into the mammalian expression vector pCAGGS and overexpressed in N2a cells. At 12 h post transfection, the cells were infected with RABV for 48 h at MOI 0.01, and virus titers in the supernatant were measured. **D.** N2a cells were transfected with pCAGGS-*Pcp4l1* (pC-*Pcp4l1*) at indicated dose for 12 h, and then infected with RABV at MOI 0.01. At 48 hpi, the virus load in the cell supernatant was measured. PCP4L1 expression level was analyzed by Western blotting. **E.** pcDNA-RABV-N, pcDNA-RABV-P together with pCAGGS or pC-Pcp4l1-flag was transfected into N2a cells for 48 h. The level of RABV-N protein and RABV-P protein was analyzed by Western blotting and normalized to GAPDH. **F.** N2a cells were transfected with pCAGGS-*Pcp4l1* (pC-*Pcp4l1*) for 12 h, and then infected with VSV at MOI 0.01. At indicated hpi, the virus load in the cell supernatant was measured. **G.** N2a cells were transfected with pC-*Pcp4l1* for 24 h, and then infected with SFV at MOI 0.01. At indicated hpi, the virus load in the cell supernatant was measured. **H.** N2a cells were transfected with pC-*Pcp4l1* for 24 h, and then infected with HSV-1 at MOI 0.01. At indicated hpi, the virus load in the cell supernatant was measured. **I.** Sequencing profile of *Pcp4l1* for ChIP-seq. The two tracks show H3K27me3 signals for pcDNA3.1 and pcDNA-EDAL samples after removing input background. The brown rectangle indicates the predicted promoter region of *Pcp4l1*. **J.** N2a cells were transfected with pcDNA-EDAL or pcDNA3.1 for 48 h, and then ChIP-qPCR were performed with H3K27me3 antibody in the promoter region of *Pcp4l1*. **K.** N2a cells were treated with 4 µM gsk126 or DMSO (mock) for 48 h and *Pcp4l1* mRNA level was analyzed by qPCR. **L.** Proposed model for EDAL-induced EZH2 lysosomal degradation, and the potential subsequent impact on EZH2-mediated epigenetic silencing of *Pcp4l1*. Statistical analysis of grouped comparisons was carried out by student’s t test(**P<0.01; ***P<0.001). Bar graph represents means ± SD, *n* = 3.

The EDAL-response genes with up-regulated transcription and the loss of H3K27me3 mark should represent candidate genes whose expression was subjected to the EDAL-EZH2 regulation, which we considered for further investigation. Six such genes were selected and evaluated whether they could restrict RABV replication. These genes were overexpressed by transient transfection in N2a cells and then RABV was infected at 12 h later. The supernatant was collect at 48 hpi and the virus titers in cell supernatant were measured. The results demonstrated that the gene encoding purkinje cell protein 4-like 1 (PCP4L1), which is small neuronal IQ motif protein closely related to the calmodulin-binding protein PCP4/PEP-19 (Bulfone, Caccioppoli et al., 2004, Morgan & Morgan, 2012), could significantly inhibit RABV replication (Fig 7C). By transfecting different amount of the plasmid expressing PCP4L1 in N2a cells, we found that PCP4L1 could inhibit RABV replication in a dose-dependent manner (Fig 7D). Furthermore, we found that PCP4L1 overexpression reduced RABV N protein level (Fig 7E), and also the virus titers of VSV, SFV and HSV-1 in N2a cells (Fig 7F-H).

ChIP-seq results showed that the H3K27me3 level on the promoter region of *Pcp4l1* was dramatically decreased after EDAL overexpression (Fig 7I), which was validated by ChIP-qPCR assay (Fig 7J). After treatment with EZH2’s inhibitor gsk126, the transcriptional level of *Pcp4l1* was significantly increased, confirming that *Pcp4l1* transcription is regulated by EZH2 (Fig 7K). All these results together suggest that EDAL might promote PCP4L1 expression by down-regulating the EZH2-mediated H3K27me3 deposition.

## DISCUSSION

We report here that multiple neurotropic viruses elicit the expression of a host lncRNA EDAL. EDAL inhibits the replication of RABV, VSV, SFV and HSV-1 in neuronal cells, and suppresses RABV infection in mouse brains. EDAL binds to the histone methyltransferase EZH2, a widely conserved epigenetic regulator, and specifically causes EZH2’s lysosomal degradation by blocking T309 *O*-GlcNAcylation. This in turn reduces cellular H3K27me3 levels. EDAL’s antiviral function resides in a 56-nt antiviral substructure that can fold into a tertiary structure with a 18-nt helix-loop that intimately contacts the T309 *O*-GlcNAcylation site of EZH2. Mutation analysis confirmed that EDAL’s effect on lysosomal EZH2 degradation requires the interaction between the 18 nt helix-loop of EDAL and EZH2 sites surrounding T309 *O*-GlcNAcylation, supporting that EDAL blocks a specific EZH2 PTM via tertiary interactions. Additionally, EDAL antiviral function could be attributed to its activated expression of a novel antiviral small peptide PCP4L1. Our discovery that neurotropicviruses elicit the expression of a neuronal antiviral lncRNA which facilitates the key epigenetic regulator EZH2 toward lysosomal degradation illustrates a way for a low level of lncRNA to effectively reduce the level of its target protein, as well as a direct biomolecular link among virus infection, host antiviral responses, and epigenetic regulation (Fig 7L). The findings of the antiviral and EZH2 degradation function carried by a 56-nt segment of EDAL and its predicted capability of folding into a functional tertiary structure together highlight a mechanism for the specificity of lncRNA actions (Fig 6C).

Recent studies have shown that post-translational modification (PTM) of EZH2 by phosphorylation affects its stability. CDK1 phosphorylates human EZH2 at T345 and T487, promoting ubiquitination of EZH2 and its subsequent degradation in proteasomes (Kaneko et al., 2010, Wu & Zhang, 2011). T345 phosphorylation site is involved in regulating EZH2 binding with HOTAIR and XIST lncRNA (Kaneko et al., 2010). K348 acetylation reduces the phosphorylation of EZH2 at T345 and T487, and increases the stability of EZH2 without interrupting PRC2 formation (Wan, Zhan et al., 2015). LncRNA ANCR facilitates the CDK1-EZH2 interaction and enhances the phosphorylation at T345 and T487, leading to EZH2 degradation and the attenuation of the invasion and metastasis of breast cancer (Li et al., 2017).

It has been recently shown that *O*-GlcNAcylation catalyzed by *O*-linked N-acetylglucosaminyltransferase (OGT) occurs at S73, S76, S84, T313, and S729 sites of the human EZH2, which does not affect the formation of the PRC2 complex. S76 and T313 are conserved in mammals, and S76A and T313A mutations independently reduce the stability of EZH2 (Chu et al., 2014, Lo et al., 2018). In the present study, molecular docking indicated that a 56-nt functional domain of EDAL lncRNA conveying both the antiviral and EZH2 degradation activity can shield T309 of mouse EZH2, the analogue of T313 in human EZH2, from the *O*-GlcNAcylation modification. PTM of biologically and therapeutically important proteins by *O*-GlcNAcylation are of interest as both lncRNA targets and as therapeutic targets. *O*-GlcNAcylation is highly abundant in eukaryotes, occurring in both the nucleus and the cytoplasm (Hanover, Krause et al., 2012, Hart, Slawson et al., 2011, Lewis & Hanover, 2014). In light of our confirmation of EDAL’s regulation of EZH2 *O*-GlcNAcylation, lncRNA regulation of other *O*-GlcNAcylation modification sites on other target regulatory (and other) proteins can be anticipated.

Note that EZH2-lncRNA interactions have been a popular model for studies of epigenetic silencing by PRC2 (Davidovich & Cech, 2015, Lee, 2012, Margueron & Reinberg, 2011, Mercer & Mattick, 2013, N, 2013, Ringrose, 2017). However, the binding specificity of PRC2 for lncRNAs and other transcripts has been challenged and re-examined recently, leading to controversy about binding specificity and promiscuity (Davidovich et al., 2015b, Davidovich et al., 2013, Wang, Goodrich et al., 2017). Our findings indicated that EDAL binds to EZH2 at a site different from that of lncRNA-HOTAIR binding of human EZH2 via residues in 342-368 region (Kaneko et al., 2010). More importantly, this study has shown that a 56-nt EDAL segment independently carries both the antiviral and EZH2 degradation function. Although we have not yet obtained structural data to support its predicted structure, our data for the function of the intimate contacts between the 18-nt helix-loop of EDAL and EZH2’s T309 *O*-GlcNAcylation site offers a new example of EZH2-lncRNA recognition and specificity.

DNA viral genome-encoded lncRNAs have recently been shown to actively interact with host epigenetic machinery to regulate both their own and host chromatin structure dynamics (Scott, 2017). Some DNA viruses repress transcription and stabilize viral latency by methylating their host’s genomic DNA (Knipe, Raja et al., 2017, Lieberman, 2016). In plants, both RNA and DNA viruses encode suppressors that limit the silencing capability of the host plants (Buchmann, Asad et al., 2009, Ruiz-Ferrer & Voinnet, 2009, Yang, Fang et al., 2013, Zhang, Chen et al., 2011). These silencing suppressors also reduce RNA-directed DNA methylation activity at transposons and repetitive sequences in the host genome, suggesting a potential regulatory role that plant viruses impose on their host epigenetic dynamics (Buchmann et al., 2009, Romanel, Silva et al., 2012, Zhang et al., 2011).

The present study reveals that neurotropic viruses elicits the expression of EDAL, a host cell lncRNA which restricts the replication of RABV, VSV, SFV and HSV-1. We experimentally link EDAL’s antiviral activity to its function in decreasing the cellular stability of EZH2, a protein whose antiviral activity has been recently revealed against the DNA virus HSV-1 (Arbuckle et al., 2017). Consequently, we found that the cellular level of H3K27me3 marks was reduced in neuronal cells, which was accompanied by the removal of in the enriched H3K27me3 mark in an antiviral gene *Pcp4l1* (Fig 7L). These findings suggest that viruses can elicit the expression of a host lncRNA which mediates EZH2 destabilization and reprograms host chromatin structure dynamics. This regulation could be anticipated during the infection by other RNA viruses and DNA virus as well. Alteration of the host epigenetic dynamics by virus-elicited host lncRNAs might not be limited to EZH2 and H3K27me3 mark. In *Drosophila*, the null mutants of the histone H3 lysine 9 methyltransferase G9a are more sensitive to RNA virus infection, and G9a controls the epigenetic state of immunity genes (Kramer, Kochinke et al., 2011, Merkling, Bronkhorst et al., 2015). It is thus possible that lncRNAs may be involved in G9a-regulated RNA virus responses.

Expression of thousands of lncRNAs has been shown to respond to DNA and RNA virus infection (Ouyang et al., 2016). Some of these lncRNAs have been shown to regulate antiviral immunity via targeting transcription factors and modulating histone modification. For example, lnc-DC binds directly to STAT3 in the cytoplasm, acting as a molecular shield to prevent STAT3 from binding to and de-phosphorylation by SHP1. As a result, lnc-DC indirectly promotes STAT3 phosphorylation on tyrosine-705 and controls human dendritic cell differentiation (Wang, Xue et al., 2014). Both mechanisms lead to the altered expression of cytokines, including IFN and TNF, as well as antiviral proteins from interferon-stimulated genes (ISGs) (Ouyang et al., 2016). It has been shown that lnc-Lsm3b binds to viral RNA sensor RIG-I as a molecular decoy, which inactive RIG-I at the late stage of viral infection and blocks type I IFN responses (Jiang, Zhang et al., 2018). Additionally, Lnczc3h7a serves as a molecular scaffold to stabilize RIG-I-TRIM25 complex and facilitates TRIM25-mediated ubiquitination of RIG-I, which promotes antiviral innate immune responses (Lin, Jiang et al., 2019). The results from EDAL in this study define epigenetic regulators as effective targets of lncRNAs in antiviral responses.

PCP4L1 is a 68 amino acids polypeptide which display sequence similarity to the Purkinje Cell Protein 4 gene (*Pcp4*) and both of which are characterized by their C-terminal IQ domain ends (Bulfone et al., 2004). PCP4L1 display a distinct expression pattern which is dominantly expressed in the CNS, and mostly expressed in circumventricular organs and modulate the production of the cerebrospinal fluid in the adult brain (Bulfone et al., 2004). Previous studies showed that PCP4L1 may be a latent calmodulin binding protein which becomes activated by post-translational modification (Morgan & Morgan, 2012). Here we demonstrate that PCP4L1 could inhibit multiple neurotropic virus infection in neuronal cells. Our results therefore reveal a novel antiviral protein which preventing the invasion of RABV, VSV, SFV, HSV-1 and maybe other neurotropic viruses into CNS.

In summary, our study of a major neurotropic virus reveals a previously unknown lncRNA-EZH2 PTM-mediated link between host antiviral responses and epigenetic regulation, and the involvement of a high specificity of lncRNA-protein tertiary interaction. The findings may reshape the current understanding of the lncRNA regulatory function, mechanism and its partnership with EZH2. EZH2 is a promising anticancer target with a well-established oncogenic role in a large variety of cancers (Conway, Healy et al., 2015, Kim & Roberts, 2016). The anticancer activities of a number of EZH2 inhibitor compounds have been reported (Kim & Roberts, 2016, Mccabe et al., 2012). The exciting finding of the 56-nt RNA substructure carrying the full EZH2 inhibitor function not only offers an example of EZH2-lncRNA recognition and specificity, but also provides new opportunity for developing anticancer and antiviral therapeutics, as well as for developing molecular tracers of EZH2 to explore the cellular activity of EZH2 during its life time.

## Materials and Methods

### Cell lines, viruses, and mice

Cell lines N2a (murine neuroblastoma N2a cells, ATCC^®^ CCL-131), BSR (a clone of BHK-21, ATCC^®^CCL-10), C8-D1A (murine astrocytes, ATCC^®^CRL-2541) and Vero (*Cercopithecus aethiops* kidney cells, ATCC^®^CCL-81) were obtained from American Type Culture Collection. BV2 (murine microglia, BNCC337749) were obtained from BeNa Culture Collection. Cells grown in a 37°C humidified 5% CO_2_ atmosphere, growth media was DMEM or RPMI1640 supplemented with 10% (vol/vol) FBS (Gibco) and 1% antibiotics (penicillin and streptomycin) (Beyotime). The recombinant rRABVs were cloned from RABV strain challenge virus standard-B2c (CVS-B2c) and constructed as described previously (Tian et al., 2016). VSV is propagated in BHK-21 cells and stored in our lab. SFV and HSV-1 is a gift from Dr. Bo Zhang (Wuhan Institute of Virology, Chinese Academy of Sciences, Wuhan, China) and Dr. Gang Cao (Huazhong Agricultural University, China), respectively, both of which are propagated in Vero cells. Female C57BL/6 mice (8 week old) mice were purchased from Hubei Center for Disease Control and Prevention, Hubei, China and housed in the animal facility at Huazhong Agricultural University in accordance with the recommendations in the Guide for the Care and Use of Laboratory Animals of Hubei Province, China. All experimental procedures involving animals were reviewed and approved by The Scientific Ethic Committee of Huazhong Agricultural University (permit No. HZAUMO-2016-009).

### Viral infection

Cells (N2a, BV2, C8-D1A and Vero) were infected with different rRABVs, VSV, SFV or HSV-1 at a multiplicity of infection (MOI) of 0.01, 0.1, 1 or 3. After 1 h at 37°C, the supernatant was discarded and cells were washed three times with PBS then cultured in DMEM or RPMI1640 supplemented with 2% (vol/vol) FBS (Gibco) and 1% antibiotics (penicillin and streptomycin, Beyotime) at 34°C in a humidified 5% CO_2_ atmosphere.

### RNA-seq library construction, sequencing and lncRNA prediction pipeline

Total RNA from RABV infected N2a cells or mock-infected cells were isolated by using Trizol^®^ reagent (Ambion) following the manufacturer’s instructions, and then treated with RQ1 DNase (Promega) to remove DNA. RNA quality and quantity were determined by measuring absorbance at 260 nm/280 nm (A260/A280) using a SmartSpec Plus spectrophotometer (BioRad). RNA integrity was verified by subjecting a sample of the RNA to electrophoresis in a 1.5% agarose gel.

Each RNA-seq library was prepared using 5 μg of total RNA. Polyadenylated mRNAs were purified and concentrated with oligo (dT)-conjugated magnetic beads (Invitrogen) and then used as templates for directional RNA-seq library preparation. Purified RNAs were iron fragmented at 95°C, followed by end repair and 5’ adaptor ligation. Reverse transcription was performed using RT primers harboring a 3’ adaptor sequence and randomized hexamer. The cDNAs were purified, amplified by PCR, and products 200–500 bp in length were isolated, quantified, and used for sequencing.

For high-throughput sequencing, the libraries were prepared following the manufacturer’s instructions and analyzed using the Illumina NextSeq500 system for 150 nt pair-end sequencing (ABlife. Inc, Wuhan, China).

### RNA-seq data processing and alignment

Raw reads containing more than two unknown (N) bases were discarded. Adaptors were removed from the remaining reads, and then short reads (less than 16 nt in length) and low quality reads (containing more than 20 low quality bases), were also excluded by using the FASTX-Toolkit sequence processing pipeline (Version 0.0.13, http://hannonlab.cshl.edu/fastx_toolkit/) to yield the final data set (clean reads). The *mus musculus* genome sequence (GRCm38) and annotation file (gencode.vM6 basic annotation) were obtained from the GENCODE database (Mudge & Harrow, 2015). Clean reads were aligned end-to-end to the mouse genome by TopHat2 (Kim, Pertea et al., 2013), allowing 2 mismatches. Reads that aligned to more than one genomic location were discarded, and uniquely localized reads were used to calculate the number of reads and RPKM values (RPKM represents reads per kilobase and per million) for each gene. Other statistics, such as gene coverage and depth, and read distribution around transcription start sites (TSSs) and transcription terminal sites (TTSs) were also obtained.

After calculating the expression levels for all genes in the samples, differentially expressed genes (DEGs) between samples were identified by edgeR (Robinson & Oshlack, 2010) using the TMM normalization method (Li, Witten et al., 2012). For each gene, the fold changes, *p-values*, and adjusted *p-values* (FDR) were also determined by the edgeR package. Genes with FDR < 0.05 were classified as DEGs.

### LncRNA prediction pipeline

The lncRNA prediction pipeline was implemented following the methods described by Liu *et al*.(Liu et al., 2017). The detailed descriptions of the prediction pipeline and filtering thresholds are as follows:

1. First, using the aligned RNA-seq data (see above), transcripts were assembled by Cufflinks V2.2.1 (Trapnell et al., 2012) using default parameters. After the initial assembly, transcripts with FPKM greater than or equal to 0.1 were subjected to a series of filters.
2. Cuffcompare (embedded in Cufflinks) was used to compare the transcripts with known mouse genes. Novel transcripts, including those that were intronic, intergenic, and antisense, were retained as candidate lncRNAs. Transcripts within 1000 bp of known coding genes were regarded as UTRs and discarded.
3. To remove potential protein-coding transcripts, coding potential score (CPS) was evaluated using the Coding Potential Calculator (CPC) (Kong, Zhang et al., 2007). CPC is a support vector machine-based classifier that assesses the protein-coding potential of transcripts based on six biologically meaningful sequence features. Transcripts with CPS scores below zero were regarded as non-coding RNAs.
4. Transcripts satisfying the above conditions, containing multiple exons and no fewer than 200 bases, or containing a single exon and no fewer than 1000 bases, were considered to be candidate lncRNAs.
5. We used Cuffmerge (from Cufflinks) to merge lncRNAs from all samples together to obtain the final lncRNA set. A total of 1662 novel lncRNA transcripts were identified, originating from 1377 lncRNA loci. The expression level of each lncRNA gene was recalculated, and antisense reads of lncRNAs were discarded.
6. Novel and known lncRNAs were combined into a single data set and subjected to analysis to identify differentially expressed lncRNA, using the same methods used to identify differentially expressed protein coding genes.

### Quantitative real-time PCR (qPCR)

Total RNA was isolated from cells and tissues by using Trizol^®^ reagent (Invitrogen). The genomic DNA was eliminated with TURBO DNA-free^TM^ Kit (Ambion, AM1907) as the manufacturer’s instructions. RNA quality was assessed by using NanoDrop 2000 (Thermo Scientific). The cDNAs were synthesized by ReverTra Ace qPCR RT Master Mix (Toyobo, FSQ-201) or First-Strand cDNA Synthesis Kit (Toyobo, FSK-101). qPCR was performed using SYBR Green Supermix (Bio-Rad). Primer sequences used in this study were listed in Appendix Table S2.

### Transfections

After seeding, cells were incubated for 12 h at 37°C. Plasmids or siRNA were transfected into cells by using Lipofectamine 3000 (Invitrogen) according to the manufacturer’s instruction.

### Rapid amplification of cloned cDNA ends (RACE)

Total RNA from N2a cells was isolated by using Trizol^®^ reagent (Invitrogen) and 5’- or 3’-RACE was performed with SMARTer^®^RACE 5’/3’ Kit (Takara, 634858) following the manufacturer’s instructions. Primers used for 5’- or 3’-RACE was designed based on the known sequence information. 5’ specific primer-GGGCTGGAGAAGTGGTTCCGTTGCTAAGGGTATTCCC; 3’ specific primer-GGGAATACCCTTAGCAACGGAACCACTTCTCCAGCC.

### Fluorescent *in situ* hybridization

The red fluorescence labeled probe (Ribo-lncRNA FISH Probe Mix) against EDAL lncRNA was designed by Ribobio Co (Guangzhou, China) and was detected by Fluorescent *In Situ* Hybridization Kit (Ribobio, R11060.1) according to the manufacturer’s instructions. Briefly, N2a cells grown on cover slips in 24-well plates were fixed with 4% (v/v) paraformaldehyde for 10 minutes (min) at room temperature then washed three times with cold PBS. And the cells were permeabilized in PBS containing 0.5％ Triton X-100 for 5 min in 4°C, then blocked in pre-hybridization buffer for 30 min at 37°C. Cells were then incubated with hybridization buffer containing probe overnight at 37°C away from light. After hybridization, cells were washed in the dark with washing buffer (4×SSC/2×SSC/1×SSC) then stained with DAPI for 10 min. Cells were again washed three times with PBS, and then imaged with an Olympus FV10 laser-scanning confocal microscope.

### EDAL specific siRNA

EDAL specific siRNA was designed and synthesized by Ribobio Co. The target sequence was 5’-GGTAGACACCCAGTGACAA-3’, and siEDAL sequence was 5‘-GGUAGACACCCAGUGACAA -3‘.

### Cell viability assay

N2a cells were transfected with plasmids, siRNAs or treated with EZH2 specific inhibitor gsk126 (Apexbio, A3446) for indicated time. The viability of N2a cells was evaluated by Cell Titer 96 AQueous One Solution cell proliferation assay kits (Promega, G3582) according to the manufacturer’s instruction.

### Construction of the recombinant RABVs (rRABV)

Mouse lncRNAs, reverse EDAL (revEDAL) were amplified from the total RNA extracted from RABV-infected N2a cells using the ReverTra Ace qPCR RT Master Mix (TOYOBO, FSQ-201) with Phanta Max Super-Fidelity DNA polymerase (Vazyme, P505-d1). The primer sets used were designed by Primer 6 (PREMIER Biosoft Biolabs) (Appendix Table S2). PCR products were digested with *BsiW*I and *Nhe*I (New England Biolabs) then ligated into the genome of recombinant RABV strain B2c (rB2c) digest used the same enzymes as previously described (Tian et al., 2016).

### Rescue of rRABVs

Recombinant RABVs were rescued as reported previously (Tian et al., 2016). Briefly, BSR cells were transfected with 2 µg of a fully infectious clone, 0.5 µg of pcDNA-N, 0.25 µg of pcDNA-P, 0.15 µg of pcDNA-G, and 0.1 µg of pcDNA-L using Lipo3000 transfection reagent (Invitrogen) according to the manufacturer’s instruction. Four days post transfection, supernatants was harvested and examined for the presence of rescued viruses using FITC-conjugated anti-RABV N antibodies (Fujirebio Diagnostics, Malvern, PA).

### Virus titration

To determine rRABV and VSV titers, BSR cells were infected with serial dilutions of the viruses. After 1 h incubation in 37°C, the cell supernatant was discarded and washed once with PBS, and then overlaid with DMEM containing 1% low melting point agarose (VWR, 2787C340). After incubation in 34°C for 72 h, the cells were stained with FITC-conjugated anti-RABV N antibody (Fujirebio Diagnostics, Malvern, PA). Then the fluorescent foci were counted under a fluorescence microscope. For VSV titration, the plaques were counted at 48 h post infection.

For SFV and HSV-1 titration, Vero cells were seeded in 12-well plates and infected with serial dilutions of the viruses. After 1 h incubation in 37°C, the cell supernatant was discarded and washed once with PBS, and then overlaid with DMEM containing 1% low melting point agarose. After incubation in 34°C for 48 h, the agarose were removed and then fixed and stained with a solution of 0.1% crystal violet and 10% formalin in PBS under UV light. After staining for 4 h, the plates were washed with water, and the plaques were counted.

### Mouse infection

Eight-week-old female C57BL/6 mice were randomly divided into indicated groups and infected intranasally with rRABV, rRABV-EDAL, rRABV-revEDAL (100 FFU) or mock infected with DMEM in a volume of 20 µl. When moribund, the mice were euthanized with CO_2_, and then the brains were collected for qPCR or immunohistochemistry analysis.

### Immunohistochemistry analysis

Groups of female C57BL/6 mice were infected intranasally with rRABV or rRABV-EDAL. At indicated times post infection (pi), mouse brains were harvested and fixed in 4% paraformaldehyde for 2 days at 4°C. Tissues were then dehydrated in 30% sucrose in PBS for 48 h at 4°C, then embedded in paraffin and sliced into 4 µm sections. For immunohistochemistry (IHC), the sections were deparaffinized and rehydrated in xylene and ethanol. Endogenous peroxidase was quenched by incubation in 3% hydrogen peroxide, and antigen retrieval was performed in 0.01 M citrate buffer. Sections were blocked then incubated with primary anti-RABV P antibody (prepared in our lab, 1:500) or CD45 antibody (Servicebio, GB11066, 1:3000) overnight at 4°C. Sections were washed again then incubated with HRP-conjugated anti-mouse (Servicebio, G1211, without dilution) or anti-rabbit secondary antibodies (Servicebio, GB23303, 1:200). After washing, sections were incubated with diaminobenzidine (ServiceBio, G1211) for color development then photographed and analyzed using an XSP-C204 microscope (CIC).

### Western blotting

N2a cells were lysed in RIPA buffer (Beyotime, P0013B) supplemented with 1x protease inhibitor cocktail (Roche). Total cell lysates were separated on 8-12% SDS-PAGE gels and transferred to PVDF membranes (Bio-Rad). Membranes were blocked with TBST with 5% (w/v) non-fat dry milk for 4 h, and probed with primary antibodies which were diluted with TBST and 5% (w/v) non-fat dry milk overnight in 4°C. The primary antibodies were against RABV N protein (prepared by our lab, 1:5000), H3K27me3 (Abclonal Technology, Wuhan, China, A2363, 1:2000), H3 (Abclonal Technology, A2348, 1:2000), EZH2 (CST, #5246, 1:2000), Flag tag (MBL, M185-3L, 1:10000), HA tag (MBL, M180-3, 1:10000), PCP4L1 (ProteinTech, 25933-1-AP, 1:2000) or GAPDH (ProteinTech, 60004-1-Ig, 1:5000). After rinsing, membranes were probed with HRP-conjugated anti-mouse (Boster, Wuan, China, BA1051) or anti-rabbit secondary antibodies (Boster, BA1055, 1:6000), then developed using BeyoECL Star kit (Beyotime, P0018A). Images were captured with an Amersham Imager 600 (GE Healthcare) imaging system.

### EDAL-EZH2 interaction 3D structure modeling

Murine EZH2 3D structure was predicted with SWISS-MODEL (https://swissmodel.expasy.org/interactive) based on human EZH2 3D structure (PDB code: 5HYN). Then amino acid sequence comparison was conducted between human EZH2 and Murine EZH2, and 98.24% similarity was calculated by Clustal2.1 (a multiple sequence alignment software, https://www.ebi.ac.uk/Tools/msa/muscle/). And the high sequence similarity ensures the authenticity of our predicted Murine EZH2 3D structure. EDAL-FD 3D structure model was predicted with RNAComposer (A automated RNA structure 3D modeling server, http://rnacomposer.ibch.poznan.pl/). In order to predict the interaction between EDAL functional domain (98-153 nt) and Murine EZH2, the template-based docking method PRIME (Zheng, Kundrotas et al., 2016) (If a template can be found, it is often more accurate than the free docking method) was used to dock the EDAL and EZH2 monomer structures at first. However, these two monomer structures could not find a suitable template in the template library, so the free docking method 3dRPC (Huang, Liu et al., 2013, Zheng, Hong et al., 2019) (A computational method was designed for 3D RNA-protein complex structure prediction.) was then utilized to dock EDAL and EZH2. Two atoms between EZH2 and EDAL with distance less than 5 angstroms in the predicted complex structure are considered to have interactions.

### RNA pull-down assay

RNA was transcribed *in vitro* with T7 RNA polymerase (Roche, 10881767001) and labeled with Biotin RNA Labeling Mix (Roche, 11685597910). The synthesized RNA was treated with Rnase-free DNase I (Thermo, EN0521) and then purified with MicroElute RNA Clean-Up Kit (OMEGA, R6247-01). The RNA was heated to 95°C for 2 min, put on ice for 5 min and then put it at room temperature for 20 min to form secondary structure. The RNA was then added to the lysed cell containing overexpressed EZH2-1-337-flag and incubated for 2 h at 4°C. Then the Streptavidin M-280 beads (Thermo Fisher Scientific, 11205D) was added to the protein-RNA mix and incubated for 1 h at room temperature. After being washed with wash buffer for three times, the samples were then analyzed by Western blotting.

### *O*-GlcNAcylation labeling and detection

The plasmid pCAGGS-EZH2-S73/S75/S725A-flag was co-tranfected with pcDNA3.1, pcDNA-EDAL or pcDNA-revEDAL in N2a cells and treated with 5 mM NH_4_Cl for 48 h. Then the cells were lysed and EZH2-S73/S75/S725A-flag was pulled down by anti-flag beads (MBL, M185-10). The extracted protein was labeled with Click-iT™ *O*-GlcNAc Enzymatic Labeling System (Invitrogen, C33368) following with the manufacture’s protocol. Then the *O*-GlcNAcylation level of the labeled EZH2-S73/S75-S725A-flag was analyzed by Click-iT™ Protein Analysis Detection Kits (Invitrogen, C33370).

### Chromatin Immunoprecipitation Sequencing (ChIP-seq) library construction and sequencing

Briefly, N2a cell were transfected with pcDNA3.1 or pcDNA-EDAL for 48 h, then the growth media of N2a cells was removed and cells were rinsed three times with cold PBS. Then cells were added with formaldehyde to a final concentration of 1% and incubated at room temperature for 10 min. To stop the cross-linking reaction, glycine was add into cells to a final concentration of 0.125 M. Cells were harvested into cold PBS by scraping and transfered into a 1.5 ml microcentrifuge tube. After centrifugation at 1000 g for 5 min at 4°C, the formaldehyde crosslinked cells were collected and resuspended in 1 ml Nuclei Lysis Buffer (50 mM Tris-HCl pH 8.0, 10 mM EDTA pH 8.0, 1 % SDS, 1 mM PMSF). Chromatin was sheared to an average size of 100-500 bp by sonication, and then centrifuged (10 min, 10000 g, 4°C). 60 µl of supernatant was diluted 10-fold with 540 µl ChIP dilution buffer (1% Triton X-100, 1.2 mM EDTA, 167 mM NaCl, and 16.7 mM Tris-HCl pH 8.0), then incubated with rotation with anti-H3K27me3 (Millipore, 07-449, 10 µg) or anti-rabbit IgG (Millipore, 12-370, 10 µg) overnight at 4°C. 50 µl protein A/G Dynabeads (Pierce™, #26162) were added to each sample and incubation continued for 2 h at 4°C on a rotating platform. Beads were pelleted then washed sequentially with low salt buffer (150 mM NaCl, 20 mM Tris–HCl pH 8.0, 0.1% SDS, 0.5% Triton X-100, and 2 mM EDTA), high salt buffer (0.1% SDS, 1% Triton X-100, 2 mM EDTA, 20 mM Tris-HCl, pH 8.1, 500 mM NaCl), LiCl buffer (0.25 M LiCl, 1% sodium deoxycholate, 10 mM Tris–HCl pH 8.0, 1% NP-40 and 1 mM EDTA), then twice with TE buffer (1 mM EDTA and 10 mM Tris–HCl pH 8.0). Chromatin was eluted from the beads by two washes with 100 µl elution buffer (100 mM NaHCO_3_, 1% SDS), the Na^+^ concentration was adjusted to 300 mM with 5 M NaCl and the crosslinks were reversed by overnight incubation in a 65°C water-bath. Samples were then incubated with 0.1 mg/ml RNase A for 1 h at 37°C, then with 1 mg/ml proteinase K for 2 h at 55°C. DNA was purified by phenol extraction and ethanol precipitation. For high-throughput sequencing, the libraries were prepared following the manufacturer’s instructions (ThruPLEX DNA-seq 48S Kit, R400427) and analyzed using an Illumina NextSeq-500 system for 150 nt pair-end sequencing (ABlife Inc., Wuhan, China).

### ChIP-seq data analysis

Adaptors and low quality bases were trimmed from raw sequencing reads using Cutadapt (Martin, 2011). Reads were aligned to the mouse-GRCm38 genome using Bowtie2 (Langmead & Salzberg, 2012). To evaluate the quality of ChIP-seq data, we performed a cross-correlation analysis, as well as FRiP and IDR analyses for the ChIP-seq data, according to the ChIP-seq guidelines provided by the ENCODE and modENCODE consortia (Kheradpour & Kellis, 2012). Peaks enriched by immunoprecipitation (compared to input DNA) were identified using MACS v1.4 (Zhang, Liu et al., 2008). We selected peaks with *p-values* less than 10^-5^. All peaks from each sample were clustered by BEDTools (Quinlan & Hall, 2010). In this step, peaks with at least 1 bp overlap or book-ended features are merged. To associate peaks with genes, we set 10000 bp as the upstream limit for the distance from the peak maximum to the TSS (transcript start site), and 3000 bp as the downstream limit for distance from the peak maximum to the TSS.

### ChIP-qPCR

Formaldehyde crosslinking of N2a cells, chromatin sonication and immunoprecipitation were performed following the same procedures as the ChIP-seq section described above. The DNA pellet was suspended in 10 µl DEPC-water. Real-time PCR was then performed using a QuantStudio 6 Flex System (ABI) according to the manufacturer’s standard protocol. Input was used to normalize the amount of each sample as an internal control. Assays were repeated at least three times and expressed as Ct values. All PCR primer sequences can be found in Appendix Table S2.

### Statistical analysis

Statistical analysis was performed using the R software (https://www.r-project.org/) or GraphPad Prism 6. Significance of differences was evaluated with either Student’s t-test, when only two groups were compared, or hypergeometric test for venn diagram. Survival percent was analyzed by log rank test. Hierarchical clustering was performed by Cluster3.0 or heatmap function in R. No statistical method was used to predetermine sample sizes. *P <0.05, **P <0.01 and ***P < 0.001.

## ACKNOWLEDGMENT

This study was partially supported by the National Program on Key Research Project of China (2016YFD0500400), the Fundamental Research Funds for the Central Universities (2662015PY227) and the National Natural Science Foundation of China (31522057). This study was also partially supported by the Appreciate the Beauty of Life Incorporation (ABL2014-09030).

## Author contributions

Conceived and designed the experiments: LZ YZ BKS. Performed the experiments: BKS DC WL QW BT JH YYL SYL JX HJ ZCL LL FH RML. Analyzed the data: BKS DC MC MZ HCC ZFF YZ LZ. Wrote the paper: BKS DC YZ LZ.

## Data availability

RNA-seq and ChIP-seq data reported in this study are deposited in Gene Expression Omnibus (GEO) with accession number GSE107310 (https://www.ncbi.nlm.nih.gov/geo/query/acc.cgi?acc=GSE107310). The data which support the findings of this study are available from the corresponding author on reasonable request.

---

**Figure EV1.**
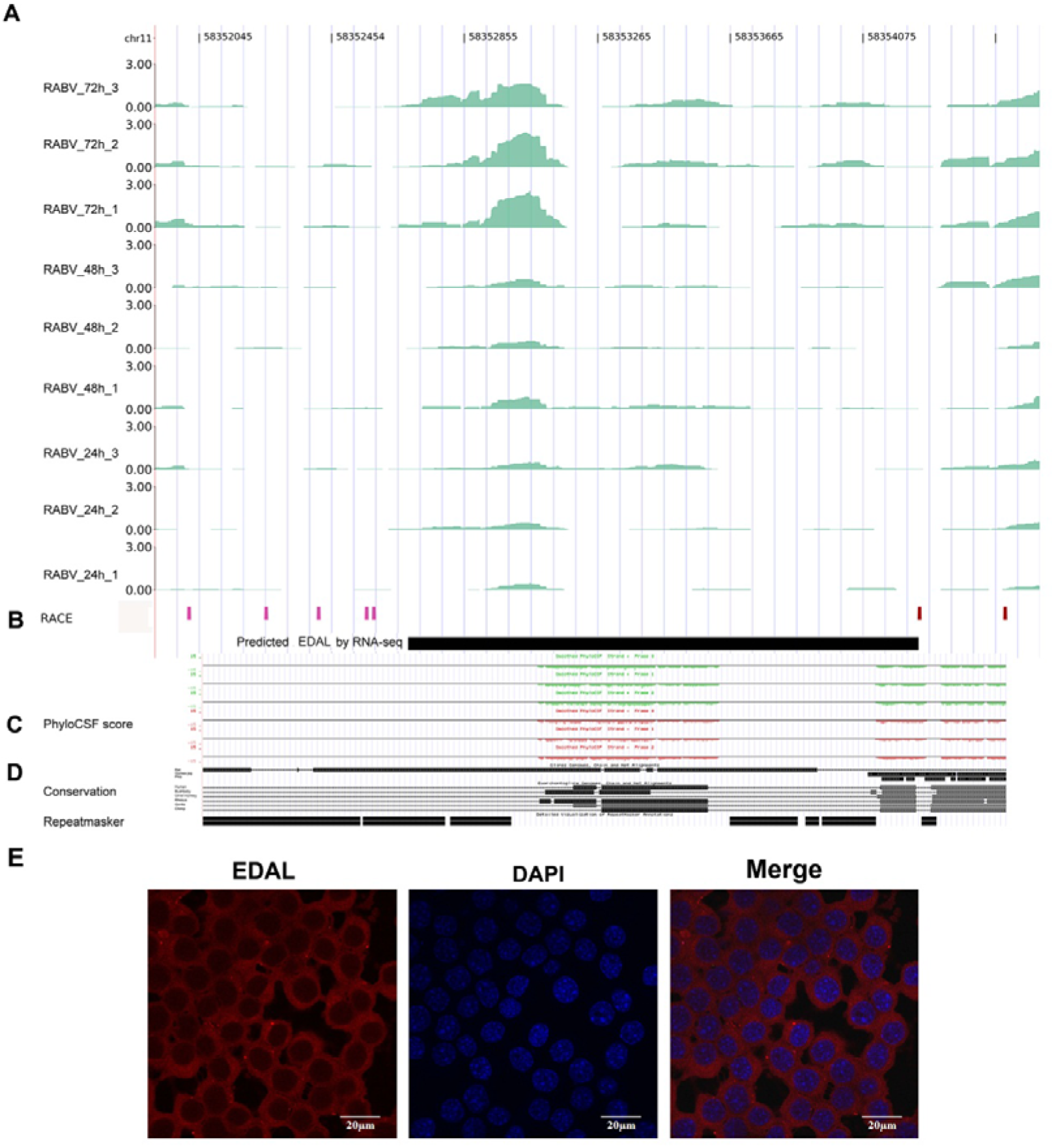
EDAL transcriptome analysis. (Related to Figure 1) **A.** Read density of EDAL. The read density is based on normalized RNA-seq signals (TPM) for each sample after RABV infection. The nine tracks show RNA-seq read density at three time points after RABV infection, with three replicates per time point. Density is shown on the y-axis. **B.** The RACE track shows the genomic location of RNA ends detected by 5’ RACE (pink) and 3’ RACE (red). The black rectangle indicates the predicted genomic location of EDAL. **C.** The PhyloCSF score track shows protein-coding scores calculated by PhyloCSF. Scores below zero indicate non-coding features. The repeated masker track shows predicted repeat sequences. **D.** Conserved and repeated sequences in EDAL. Sequence analyses were performed using the UCSC genome browser. **E.** RNA fluorescent *in situ* hybridization (FISH) assay were performed in N2a cell. Red-EDAL, Blue-4’, 6-Diamidino-2-phenylindole dihydrochloride (DAPI). Scale bar, 20 μm.

**Figure EV2.**
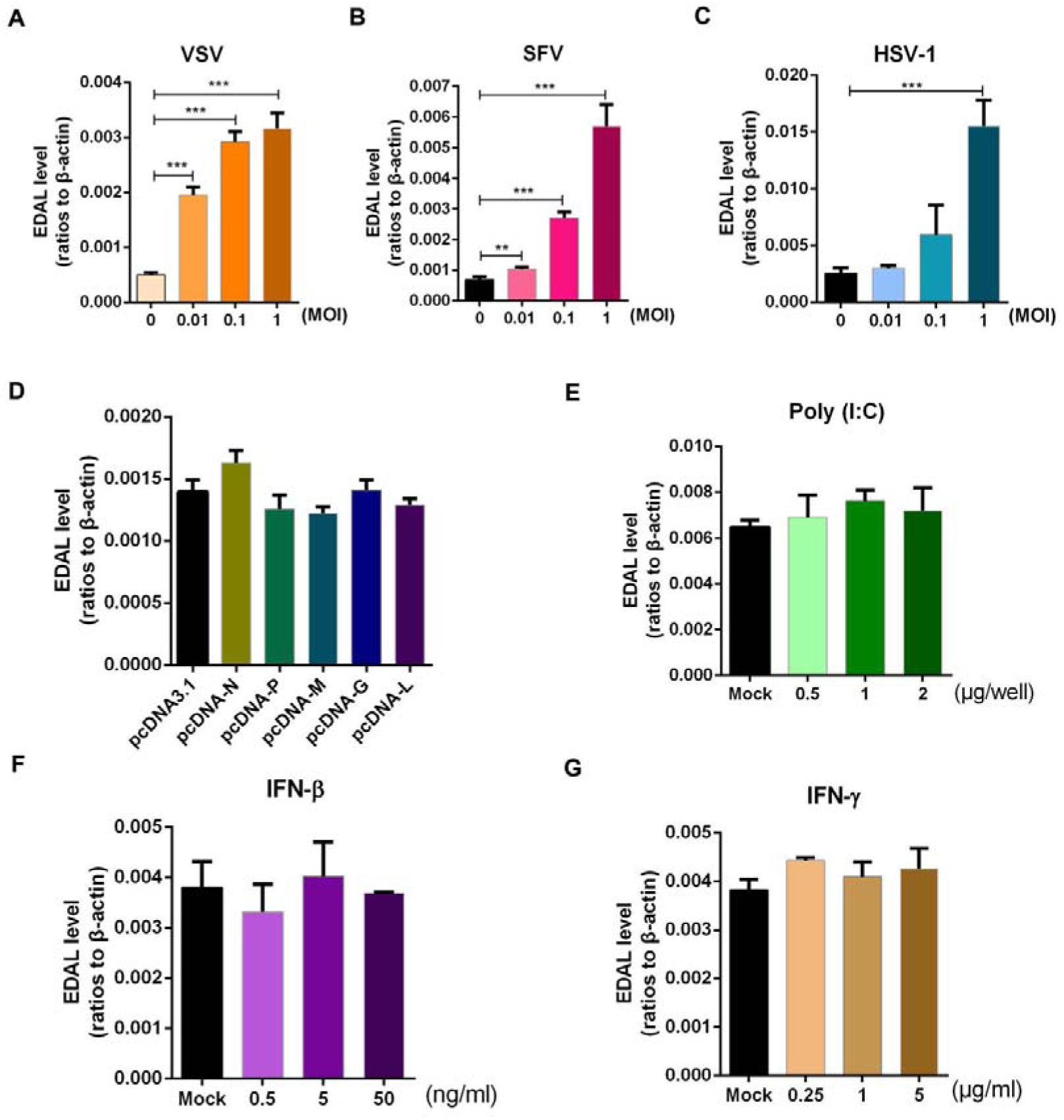
EDAL is not up-regulated by RABV proteins, dsRNA, or interferons. (Related to Figure 1) **A.** N2a cells were infected with VSV at different MOIs for 12 h and EDAL level was analyzed by qPCR. **B.** N2a cells were infected with SFV at different MOIs for 18 h and EDAL level was analyzed by qPCR. **C.** N2a cells were infected with HSV-1 at different MOIs for 18 h and EDAL level was analyzed by qPCR. **D.** N2a cells were transfected with plasmids expressing different RABV proteins. EDAL levels were analyzed by qPCR at 24 h post transfection. **E.** N2a cells were transfected with poly(I:C) (a mimic of dsRNA) at indicated doses. EDAL levels were measured by qPCR at 24 h post transfection. **F,G** N2a cells were treated with IFN-β (**F**) or IFN-γ (**G**) for 24 h. EDAL levels were analyzed by qPCR. Statistical analysis of grouped comparisons was carried out by student’s t test(**P<0.01; ***P<0.001). Bar graph represents means ± SD, *n* = 3.

**Figure EV3.**
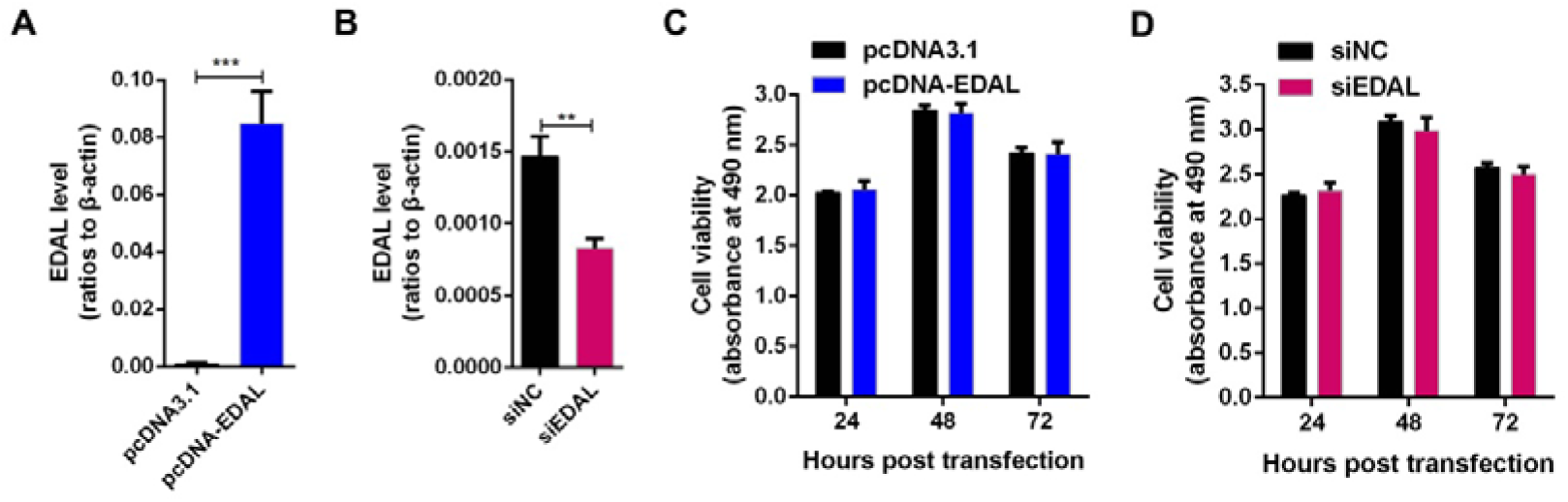
Cell viability post overexpressing or silencing EDAL. (Related to Figure 2) **A.** EDAL was cloned into a mammalian expression vector pcDNA3.1, named pcDNA-EDAL. After transfection in N2a cells, the expression level of EDAL was measured by qPCR. **B.** N2a cells were transfected with EDAL specific siRNA (siEDAL) or siNC then the level of EDAL was confirmed by qPCR. **C.** N2a cells were transfected with pcDNA3.1 or pcDNA-EDAL for indicated times, cell viability was evaluated using a Cell Titer 96 AQueous One Solution cell proliferation assay kits (G3582) from Promega. **D.** N2a cells were transfected with siEDAL or siNC for indicated times, cell viability was measured. Statistical analysis of grouped comparisons was carried out by student’s t test(\**P* < 0.05; **P<0.01; ***P<0.001). Bar graph represents means ± SD, *n* = 3.

**Figure EV4.**
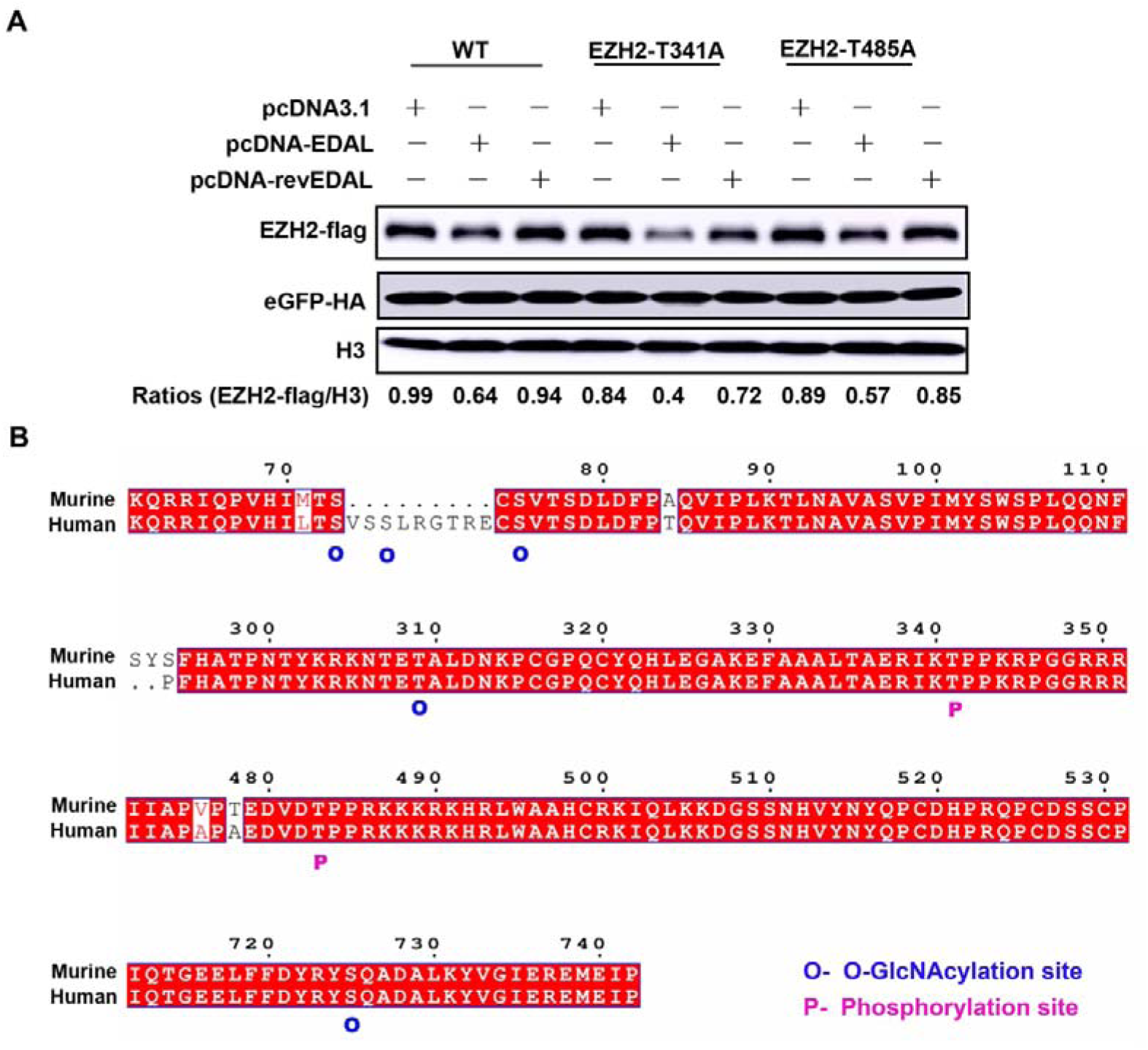
Amino acid sequence comparison between murine and human EZH2. (Related to Figure 6) **A.** The potential phosphorylation sites of murine EZH2 was mutated into A. Then the mutated EZH2 was expressed together with pcDNA3.1, pcDNA-EDAL or pcDNA-revEDAL in N2a cells for 48 h. Then EZH2-flag level was analyzed by Western blotting and normalized to H3. **B.** The amino acid sequence of murine and human EZH2 were aligned by using an online software ESPript3.0 (http://espript.ibcp.fr/ESPript/cgi-bin/ESPript.cgi). The *O*-GlcNAcylation sites and phosphorylation sites of human EZH2 were marked by O (*O*-GlcNAcylation) or P (phosphorylation), respectively.

## Supplementary Information

### Supplementary Figures

**Appendix Figure S1.**
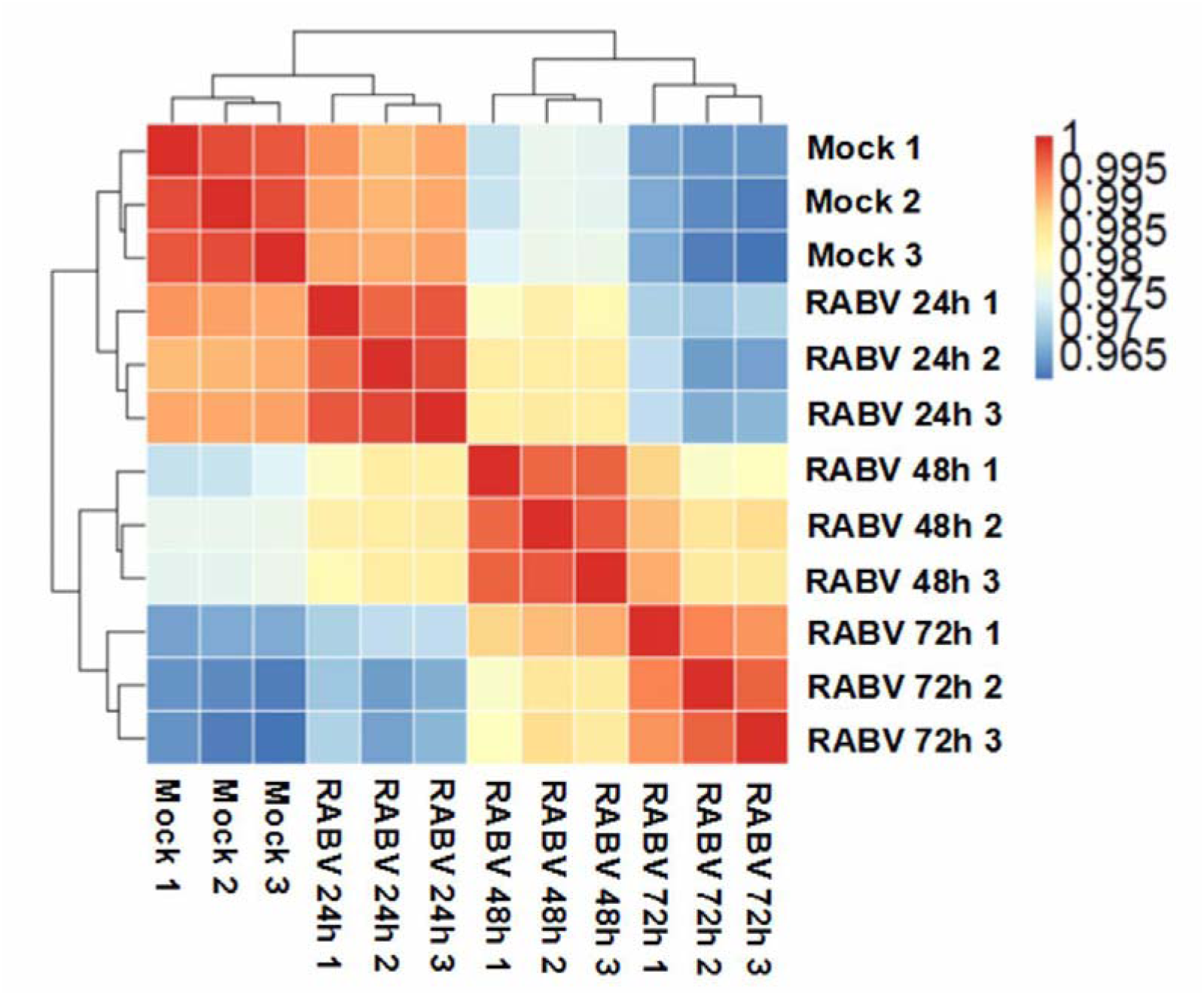
Sample correlation analysis. Hierarchical clustering heatmap shows global transcriptional changes after RABV infection. The Pearson correlation coefficients (PCCs) for each sample pair are represented using the colors in the color bar to indicate coefficient magnitude.

**Appendix Figure S2.**
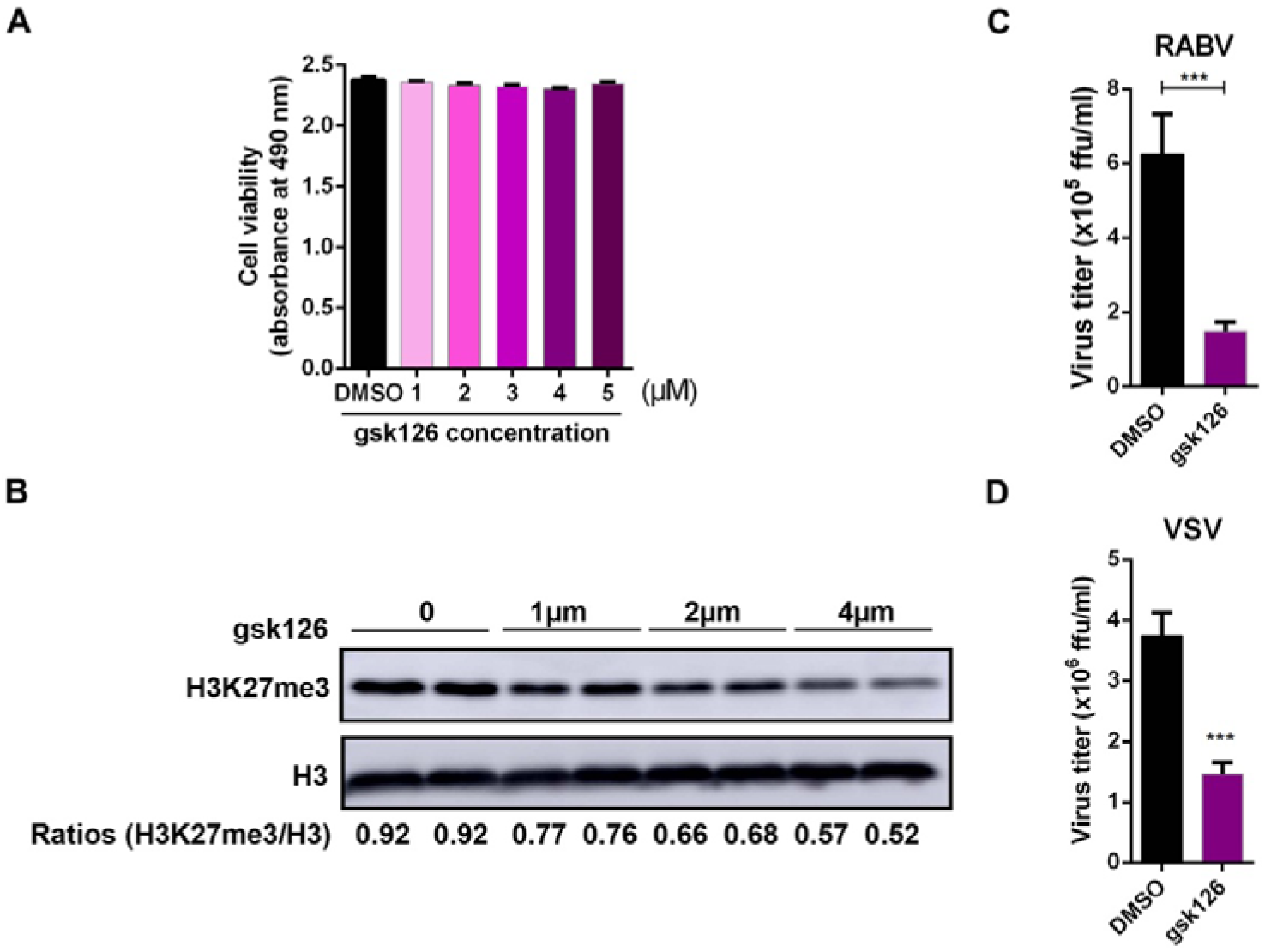
EZH2 specific inhibitor gsk126 inhibits RABV and VSV replication in N2a cells. **A,B.** After treatment with different concentrations of gsk126, an EZH2 specific inhibitor, the viability of N2a cells was evaluated by using Cell Titer 96 AQueous One Solution cell proliferation assay kit (Promega, Madison, WI) (**A**). (n=3) H3K27me3 levels were measured by Western blotting and normalized to H3 (**B**). **C.** N2a cells were treated with 4 µM gsk126 or DMSO for 12 h, and then infected with rRABV at MOI 0.01. At 48 hpi., the virus load in the supernatant was titrated. **D.** N2a cells were treated with 4 µM gsk126 or DMSO for 12 h, then infected with VSV at MOI 0.01 for 12 h, the virus load in the supernatant were measured. Statistical analysis of grouped comparisons was carried out by student’s t test(***P<0.001). Bar graph represents means ± SD, *n* = 3.

**Appendix Table S1.** Sequencing and mapping information of ChIP-seq experiments. Each sample was tested in duplicates.

| Sample | Raw reads | Reads after QC | total mapped reads | uniquely mapped reads | multiple mapped reads |
| --- | --- | --- | --- | --- | --- |
| pcDNA3.1_H<br>3K27me3_1 | 6993285<br>8 | 55668777 | 52437993<br>(94.20%) | 39942525<br>(76.17%) | 12495468<br>(23.83%) |
| pcDNA3.1_H<br>3K27me3_2 | 6789127<br>8 | 45142907 | 42187626<br>(93.45%) | 30301699<br>(71.83%) | 11885927<br>(28.17%) |
| pcDNA3.1_in<br>put_1 | 7899434<br>0 | 65958160 | 63686273<br>(96.56%) | 46843529<br>(73.55%) | 16842744<br>(26.45%) |
| pcDNA3.1_in<br>put_2 | 7631897<br>6 | 55917681 | 53343077<br>(95.40%) | 38161225<br>(71.54%) | 15181852<br>(28.46%) |
| pcDNA-EDAL<br>_H3K27me3_<br>1 | 7394171<br>2 | 58906209 | 55816652<br>(94.76%) | 40379662<br>(72.34%) | 15436990<br>(27.66%) |
| pcDNA-EDAL<br>_H3K27me3_<br>2 | 6808576<br>8 | 45952519 | 42358792<br>(92.18%) | 31024033<br>(73.24%) | 11334759<br>(26.76%) |
| pcDNA-EDAL<br>_input_1 | 7233516<br>4 | 59681813 | 57346675<br>(96.09%) | 41376294<br>(72.15%) | 15970381<br>(27.85%) |
| pcDNA-EDAL<br>_input_2 | 7056582<br>4 | 51660023 | 49080890<br>(95.01%) | 35607692<br>(72.55%) | 13473198<br>(27.45%) |

**Appendix Table S2.**
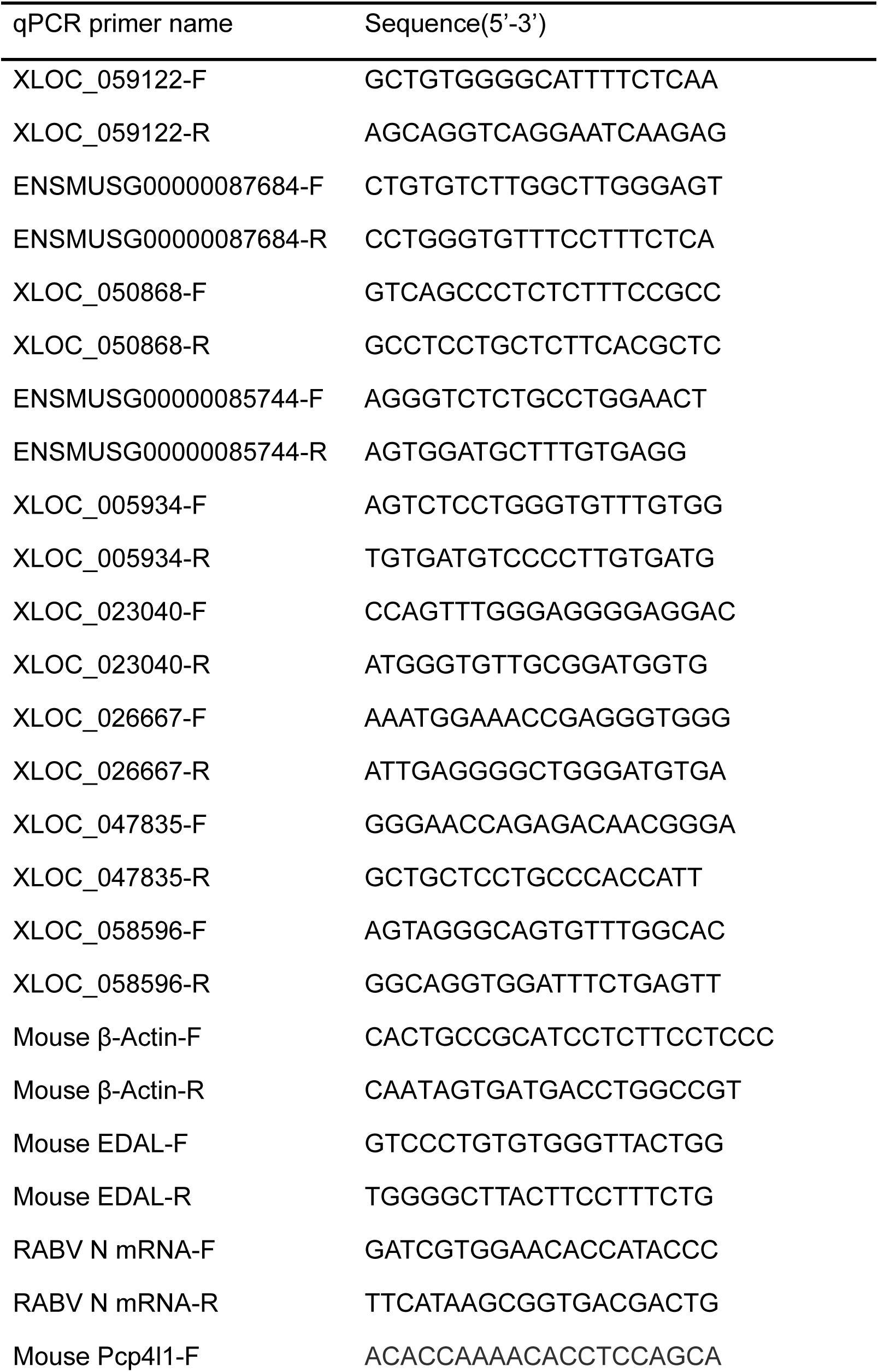
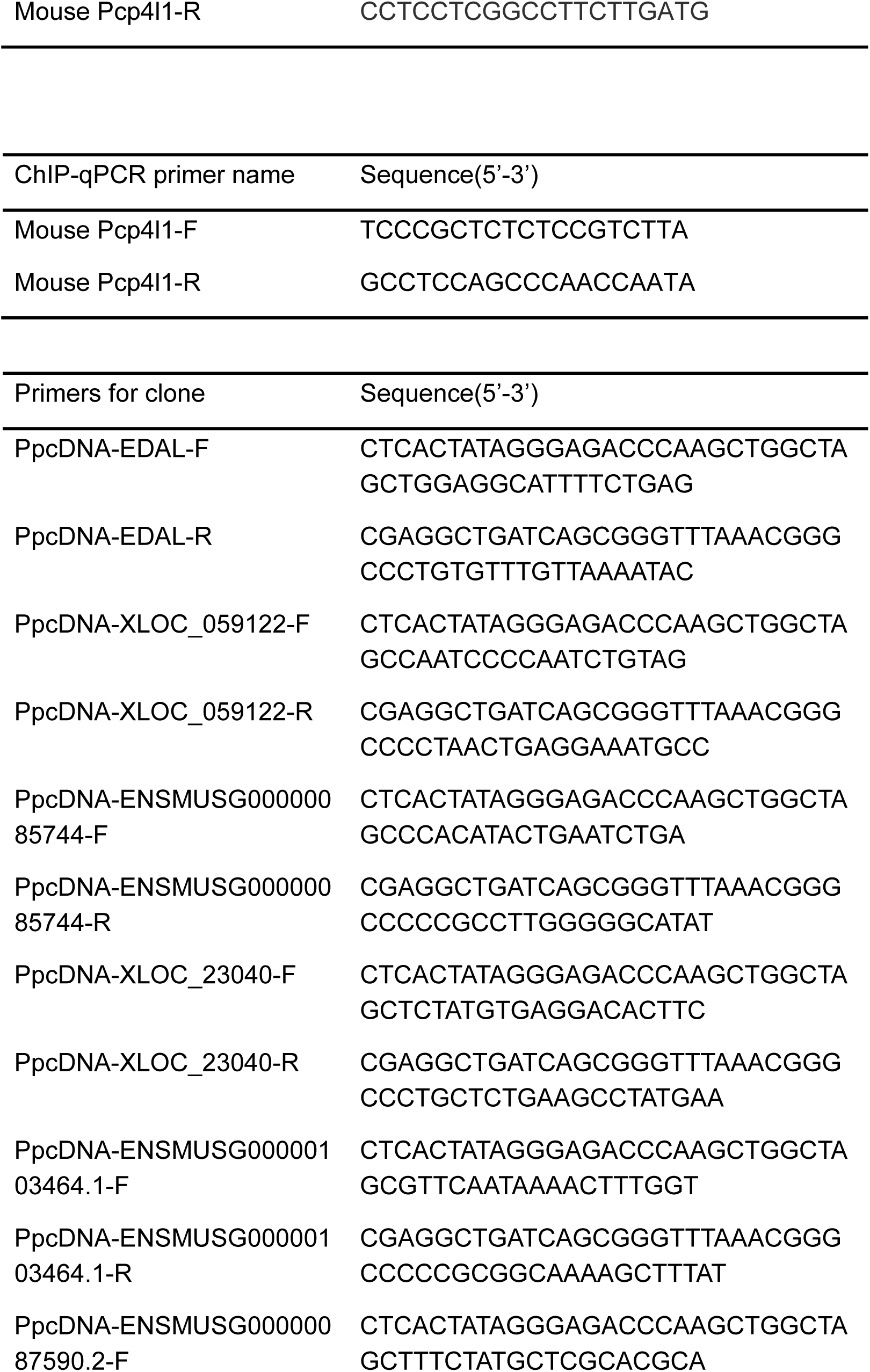
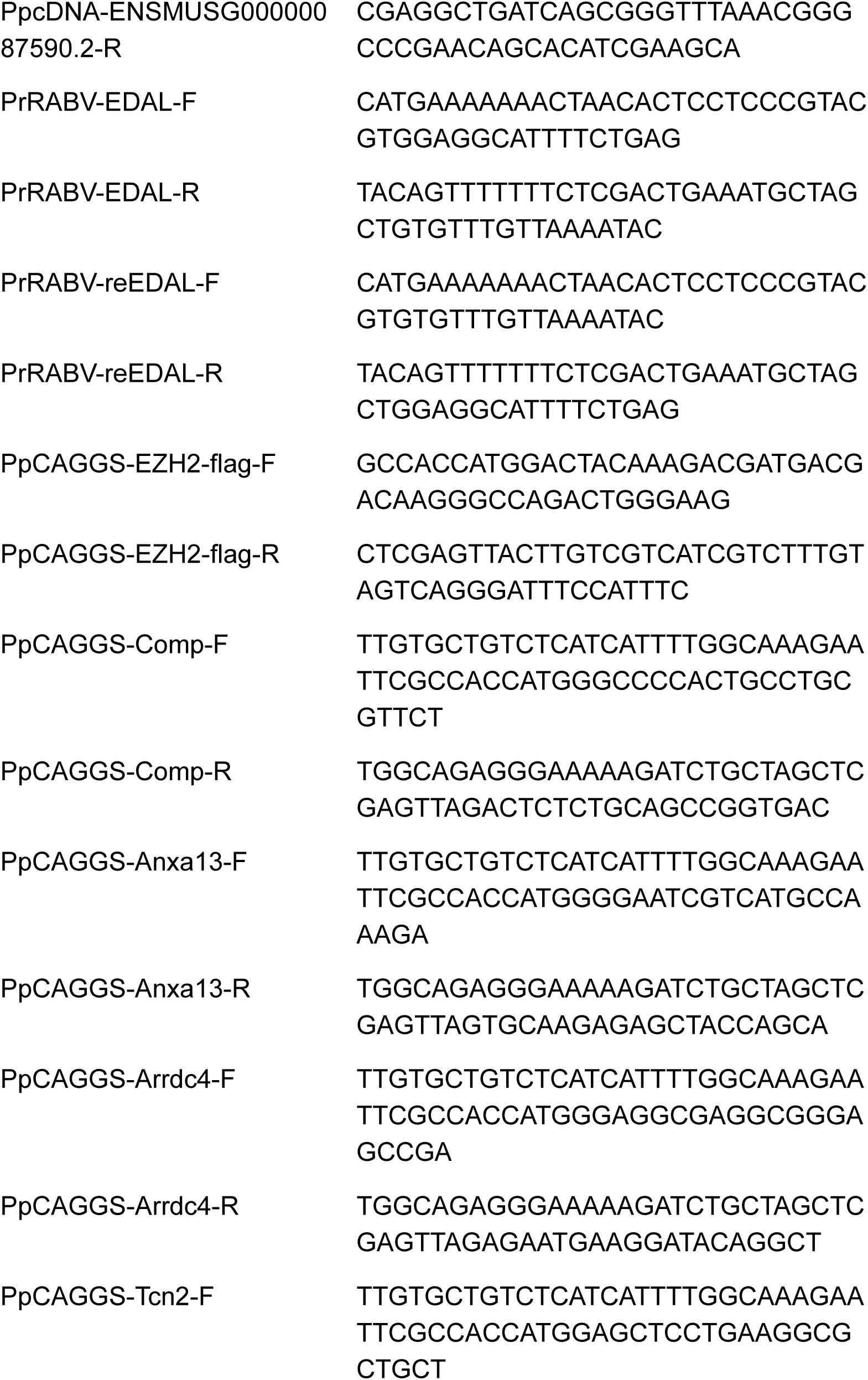
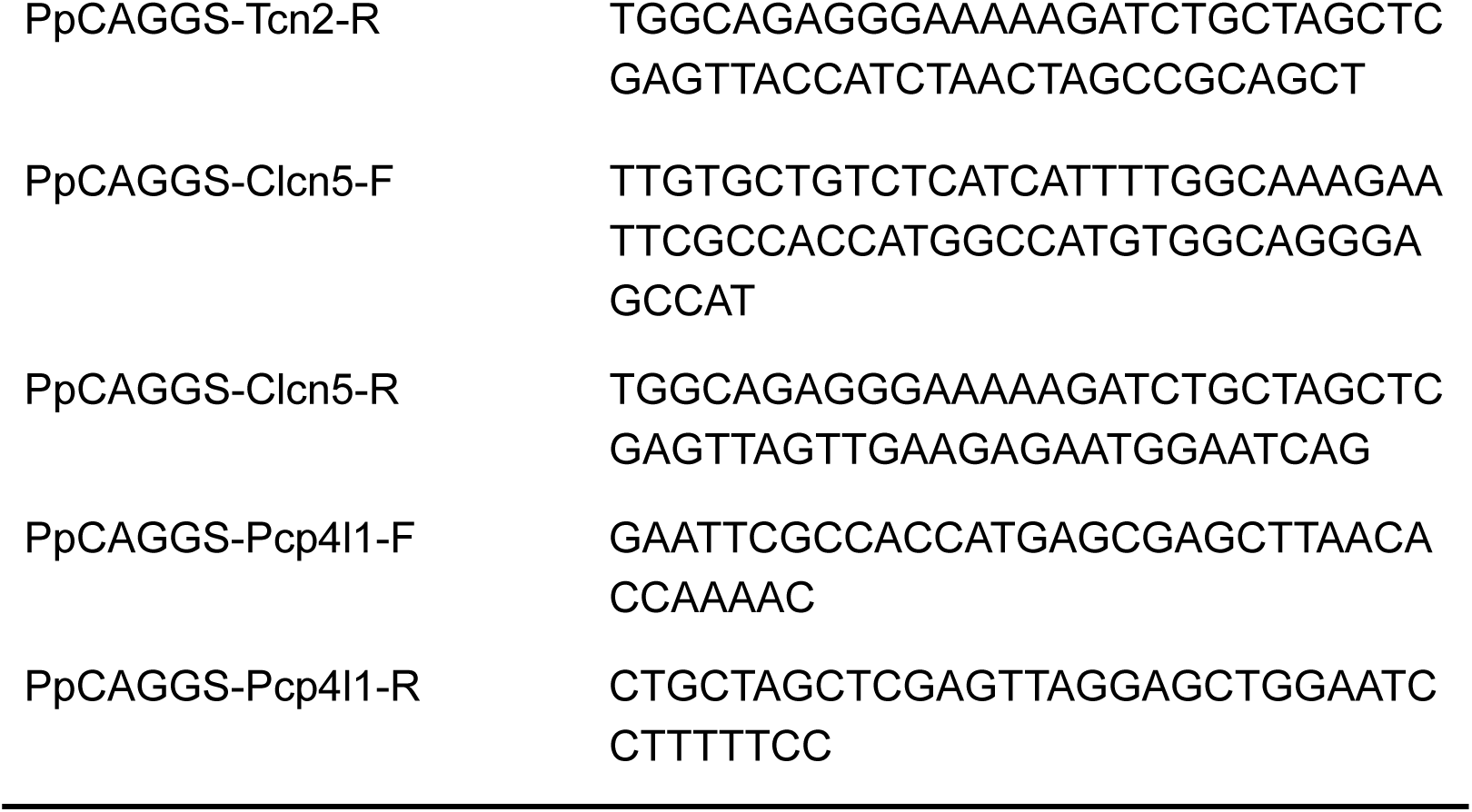
The primer sets used in this study.

